# Molecular Rewiring in TNNT2-Linked Hypertrophic Cardiomyopathy hiPSC-Derived Cardiomyocytes upon Metabolic Maturation

**DOI:** 10.1101/2025.09.14.676131

**Authors:** Homa Hamledari, Farah Jayousi, Sanam Shafaattalab, Saif Dababneh, Ramon Klein Geltink, Philipp Lange, Maksymilian Prondzynski, Glen Tibbits, Sheila Teves

## Abstract

Human induced pluripotent stem cell–derived cardiomyocytes (hiPSC-CMs) are widely used to model cardiac development and inherited cardiomyopathies, yet their immature metabolic state limits interpretation of disease-associated molecular programs. While multiple strategies promote structural and functional maturation, less is known about the molecular regulation of metabolic maturation as a distinct developmental transition. Here, we examine how metabolic maturation reshapes molecular and metabolic states in wild-type and *TNNT2*-linked hypertrophic cardiomyopathy (HCM) hiPSC-CMs. Using integrated transcriptomic, chromatin accessibility, proteomic, and metabolic profiling, we define the molecular trajectory associated with metabolic maturation in wild-type hiPSC-CMs, characterized by coordinated transcriptional and epigenetic remodeling, enhanced mitochondrial oxidative metabolism, and progressive suppression of mTORC1 signaling. In contrast, hiPSC-CMs carrying *TNNT2* HCM variants (I79N^+/−^ and R278C^+/−^) exhibit variant-specific deviations from this metabolic maturation trajectory. While early CM differentiation is largely preserved, metabolic maturation reveals defects in mitochondrial respiration and chromatin organization, with the more clinically severe I79N^+/−^ variant showing sustained metabolic impairment. We further find that mTORC1 activity is temporally misregulated during metabolic maturation in HCM hiPSC-CMs, with reduced signaling at early stages and normalization at later time points. Pharmacological inhibition of mTORC1 with rapamycin partially improves disease-associated protein expression signatures in the I79N^+/−^ variant, particularly when applied during early differentiation. Together, these findings demonstrate that TNNT2-linked HCM involves disrupted metabolic maturation programs coupled to altered gene regulatory states, highlighting metabolic maturation as a critical context for studying cardiomyopathy-associated molecular phenotypes in hiPSC-CMs.

## INTRODUCTION

Cardiovascular diseases remain the leading cause of global mortality, underscoring the need for human-relevant models that accurately capture both cardiac development and disease mechanisms^1^. Human induced pluripotent stem cell-derived cardiomyocytes (hiPSC-CMs) have emerged as a powerful platform for modeling inherited cardiomyopathies and testing therapeutic strategies ^2^. However, a major limitation of hiPSC-CMs is their immature phenotype, which more closely resembles fetal than adult CMs ^3^. While structural and electrophysiological immaturity of hiPSC-CMs has been widely recognized, metabolic maturation, defined by the transition from glycolytic to oxidative energy production, represents a distinct developmental program that remains understudied ^3^.

To promote hiPSC-CM maturation, a wide range of strategies has been developed, including electrical and mechanical stimulation, three-dimensional culture systems, and metabolic or hormonal interventions ^4–8^. In particular, metabolic maturation approaches that alter substrate availability and hormonal signaling, such as fatty acid supplementation, low glucose, thyroid hormone, glucocorticoids, and growth factor modulation, have been used to promote metabolic remodeling in hiPSC-CMs ^9–13^. These interventions robustly induce changes in mitochondrial substrate utilization and energy metabolism, even in the absence of complete structural or electrophysiological maturation, making them a useful system for interrogating metabolic state transitions in isolation ^4^. Importantly, the molecular trajectories underlying metabolic maturation, particularly at the level of gene regulation and chromatin organization, remain incompletely defined.

Most molecular studies of hiPSC-CM differentiation have focused on early lineage specification, with less attention paid to the metabolic maturation phase as a distinct and regulated developmental transition ^14–18^. Emerging evidence suggests that metabolic remodeling during maturation is tightly coupled to epigenetic and transcriptional reprogramming, raising the possibility that disease-associated phenotypes may only become apparent once CMs engage adult-like metabolic programs ^19,20^. This is especially relevant for inherited cardiomyopathies, where pathogenic variants may perturb the ability of CMs to adapt to increasing energetic and mechanical demands during maturation ^21^.

Hypertrophic cardiomyopathy (HCM) is a common inherited cardiac disorder most often caused by variants in sarcomeric proteins ^22^. Specifically, variants in cardiac troponin T (TNNT2) account for a subset of HCM cases and display marked heterogeneity in clinical severity, age of onset, and penetrance ^23,24^. The *TNNT2* variants I79N^+/−^ and R278C^+/−^ exemplify this spectrum, with I79N^+/−^ associated with more severe disease and R278C^+/−^ representing a more frequent, intermediate-effect variant ^25,26^. These sarcomeric HCM variants increase energetic demand through altered contractile dynamics ^27,28^, raising the possibility that disease-associated phenotypes may emerge specifically during metabolic state transitions rather than early differentiation. Studying metabolic maturation independently of full structural maturation therefore provides a controlled context to uncover disease-relevant metabolic and gene regulatory changes.

One pathway that integrates metabolic status, cellular growth, and stress adaptation is the mechanistic target of rapamycin complex 1 (mTORC1)^29^. Activity of mTORC1 is sensitive to nutrient availability and has been implicated in both CM maturation and cardiomyopathy pathogenesis^30–34^. While mTOR signaling has been studied in the context of several sarcomeric HCM variants^35,36^, its temporal regulation during CM maturation, particularly in *TNNT2*-linked HCM, remains poorly understood. In this study, we focus specifically on metabolic maturation as a defined developmental transition in hiPSC-CMs. Using genome-edited *TNNT2* I79N^+/−^ and R278C^+/−^ hiPSC-CMs and isogenic controls, we examine how metabolic maturation reshapes transcriptional, epigenetic, proteomic, and metabolic states, without asserting global structural or functional maturity. By comparing wild-type and HCM-specific metabolic maturation trajectories, we identify variant- and stage-dependent defects and uncover temporal misregulation of mTORC1 signaling. These findings establish metabolic maturation as a distinct and informative context for investigating cardiomyopathy-associated molecular phenotypes in stem cell-based models.

## Materials and methods

### hiPSC culture and cardiac monolayer differentiation

Our control hiPSC cell line (ID: iPS IMR90-1; WiCell Research Institute) was genome-edited to introduce TNNT2 I79N and R278C variants, as described previously^25,37^. For routine hiPSCs culture cells were seeded at a density of 200,000 cells per well in Matrigel-coated plates (0.5 mg per 6-well plate, Thermo Fisher Scientific, #354277). hiPSCs were maintained in mTeSR Plus medium (Stem Cell Technologies, #100-0276) and passaged every 4 days using ReLeSR (Stem Cell Technologies, #100-0483). At day 0 of differentiation (∼70-80% confluency), the mTeSR medium was replaced with RPMI 1640 medium (Thermo Fisher Scientific, # 11875093) supplemented with 2% B27 without insulin (Thermo Fisher Scientific, #A1895601) and 12 µM CHIR99021 (R&D Systems, # 4953) for 24 hours, after which the medium was changed. On day 3 of differentiation, 5 µM IWP4 (Tocris, #5214) was added without a medium change until the subsequent medium change on day 5. On day 7 of differentiation, RPMI 1640 medium was supplemented with B27 with insulin (Thermo Fisher Scientific, #17504044). Beating monolayers were observed from days 9-12, and the medium was changed every 3 days thereafter. Cells were maintained in RPMI 1640 + B27 with insulin for cardiomyocyte maintenance. For metabolic maturation treatments, hiPSC-CMs were treated with maturation media including 1 µM T112 (Tocris, #7690), 1 µM Dexamethasone (Sigma, #D4902), 11 nM IGF1 (StemCell Technologies, #78142), 4 nM T3 (Sigma, #T6397), complementing low glucose (3 mM, Sigma, #G7021) and fatty acid-enriched media as described by^9^.

### Drug treatment

10 nM Rapamycin (Millipore Sigma, #55-321) was applied for 2 weeks, with fresh medium changed every 3 days.

### Gene expression analysis

RNA was extracted from hiPSC-CMs at day 30 and 44 of differentiation using the TRIzol method (Invitrogen, #15596026) following the manufacturer recommendations. The Agilent 2100 Bioanalyzer was used for RNA quality assessment, and samples with RIN values above 9 were submitted for RNA sequencing. The Illumina Stranded mRNA Prep (Illumina) protocol was used for sample processing. Sequencing was carried out on the Illumina NextSeq 2000 platform with paired-end 59 bp × 59 bp reads. The sequencing reads were aligned to the *Homo sapiens* hg38 reference genome using the DRAGEN RNA pipeline version 4.3 (Illumina). DESeq2 differential gene expression analysis was performed, and functional enrichment of significant genes (adjusted p-value < 0.05, absolute fold change ≥ 1.5) was carried out using Gene Ontology (GO) enrichment analysis. Visualizations, including PCA, volcano plots, gene expression, and correlation heatmaps, were generated in R (version 4.2.3).

### Cleavage Under Targeted Accessible Chromatin (CUTAC)

Chromatin accessibility profiling was performed according to a study by Henikoff et al^38^. To extract cardiomyocytes, diluted trypsin (0.05%) was used with 50,000 cardiomyocytes per replicate. Ten percent Drosophila S2 cells were added during the primary washing step for spike-in normalization. Antibody incubation was performed using RNA Pol II S5P (Cell Signaling, #D9N5I), rabbit IgG (Cell Signaling, #2729), H3K4me2 (Epicypher, #13–0027), and guinea pig anti-rabbit IgG secondary antibody (Antibodies-Online, #ABIN101961). The pAG-Tn5 enzyme (Epicypher, #15-1017) was used and activated to cleave accessible chromatin regions surrounding RNA Pol II binding sites. Twenty percent N,N-dimethylformamide (Sigma-Aldrich, #D-8654) was included in the tagmentation buffer of the CUTAC protocol. Subsequently, PCR was used to prepare sequencing libraries, and samples were quality checked using the TapeStation (Agilent Technologies, Inc) before pooling for sequencing on the NextSeq 2000 with a 50-cycle kit.

### Statistics and data visualization

GraphPad Prism (version 9.5.1) was used for data visualization. One-way ANOVA was used to compare multiple groups. For comparisons between two groups, t-tests with False Discovery Rate (FDR) correction were applied. A p-value < 0.05 was considered statistically significant.

**Additional methods can be found in Supplementary Information.**

## RESULTS

### Maturation-inducing factors promote a shift toward oxidative metabolism and fatty acid utilization

During postnatal CM development, cellular energy production shifts from glycolysis toward mitochondrial oxidative phosphorylation, accompanied by increased reliance on fatty acids as a fuel source^39^. To model this aspect of CM metabolic maturation, we differentiated human iPSCs into CMs (hiPSC-CMs) using established stage-specific WNT modulation protocols, followed by two rounds of metabolic selection with sodium L-lactate to enrich for CMs ^9,40^. After differentiation, hiPSC-CMs were treated for four weeks with a fatty acid–enriched maturation medium (FFA MM), plus an estrogen-related receptor agonist (T112), a combination of thyroid hormone (T3), dexamethasone (DEX) insulin-like growth factor 1 (IGF1), or with the all five components together (maturation-inducing factor, MIF), as outlined in the experimental timeline (**Figure 1A**). Control samples were treated with differentiation media (i.e., RPMI1640), keeping them in immature state, with pronounced differences in morphological features between control and MIF-treated samples (**Figure S1A**).

**Figure 1.**
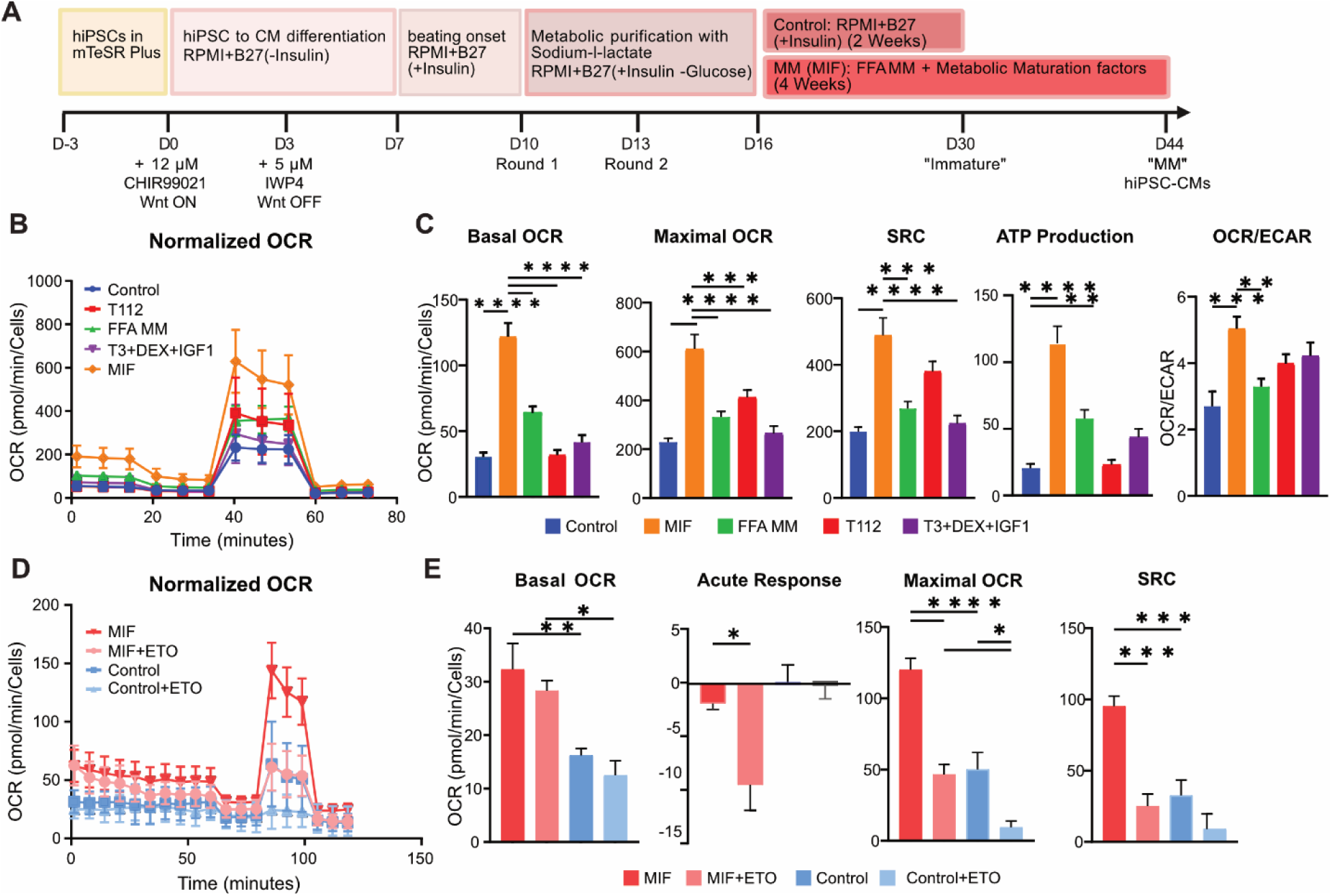
Metabolic profiling of MIF-treated hiPSC-CMs using Seahorse Mito Stress Test analysis. **(A)** Schematic of the differentiation and maturation timeline. **(B)** Oxygen consumption rate (OCR) kinetics normalized to cell number in RPMI-treated controls, individual maturation-inducing factor treatments, and the combined maturation-inducing factor (MIF) condition. **(C)** Quantification of basal respiration, maximal respiration, spare respiratory capacity (SRC), and OCR/ECAR ratio (*n*= 7-9 independent differentiations). **(D)** Fatty acid oxidation (FAO) stress test showing OCR kinetics in MIF-treated hiPSC-CMs. **(E)** Quantification of basal OC, and Basal following etomoxir (ETO) treatment (i.e., Acute response), maximal OCR, and SRC during FAO stress testing. Statistical significance was determined using one-way ANOVA with Tukey’s post hoc test (*n* = 5-6 independent differentiations). *\*p ≤ 0.05, **p ≤ 0.01, ***p ≤ 0.001, ****p ≤ 0.0001*.

To assess metabolic maturation, we performed Seahorse mitochondrial stress tests on Day 44 and quantified mitochondrial oxygen consumption rate (OCR) as a readout of oxidative metabolism. Treatment with the combined MIFs resulted in a pronounced increase in basal, maximal respiration, and ATP production compared to control conditions or individual factors alone (**Figure 1B–C**). Notably, spare respiratory capacity (SRC), reflecting the ability of cells to respond to increased energetic demand, was the most significantly enhanced parameter in MIF-treated hiPSC-CMs (**Figure 1C**). Consistent with a shift toward oxidative metabolism, MIF-treated cells also exhibited a marked increase in the basal OCR-to-extracellular acidification rate (ECAR) ratio, indicative of reduced reliance on glycolysis relative to mitochondrial respiration (**Figure 1C, Figure S1B**).

To directly examine fatty acid utilization, we inhibited mitochondrial fatty acid β-oxidation using etomoxir (ETO), which has been shown to delay mouse CM maturation ^41^. In MIF-treated hiPSC-CMs, ETO treatment significantly reduced basal OCR (i.e., acute response), whereas control hiPSC-CMs showed no significant change (**Figure 1D,E**). Maximal respiration was reduced under both conditions, but the effect was more pronounced in MIF-treated cells (**Figure 1E**). In contrast, respiratory reserve was largely unaffected by ETO in control hiPSC-CMs (**Figure 1E**), indicating limited dependence on fatty acid oxidation in the absence of MIF treatment.

Together, these data demonstrate that combined MIF treatment induces a metabolic shift toward increased mitochondrial oxidative metabolism and fatty acid utilization, consistent with metabolic maturation of hiPSC-CMs. Based on these findings, we used the combined MIF condition in subsequent experiments to investigate how metabolic maturation reshapes molecular and regulatory landscapes in wild-type and TNNT2-linked HCM hiPSC-CMs.

### MIF treatment induces coordinated transcriptional, epigenetic, and proteomic remodeling associated with metabolic maturation

To define the molecular programs accompanying MIF-induced metabolic maturation, we performed integrated transcriptomic, chromatin accessibility, and proteomic profiling of control and MIF-treated hiPSC-CMs. Across all three modalities, MIF treatment elicited coordinated and widespread remodeling consistent with a shift toward oxidative metabolism and fatty acid utilization.

We first performed bulk mRNA sequencing on control and MIF-treated hiPSC-CMs (three biological replicates per condition). Principal component analysis (PCA) and pairwise correlation revealed high reproducibility between replicates and clear segregation between conditions (**Figure S2A**). Differential expression analysis identified 3,132 upregulated and 1,258 downregulated genes following MIF treatment (adjusted *p* < 0.05, |log₂FC| > 1.5; **Figure 2A**). Transcriptional changes prominently reflected metabolic remodeling, with increased expression of genes involved in mitochondrial oxidative metabolism and lipid handling (including ACADL, CKMT1B, and PDK4) and concomitant downregulation of glycolysis-associated genes (PFKP, HK2, PGAM2).

**Figure 2.**
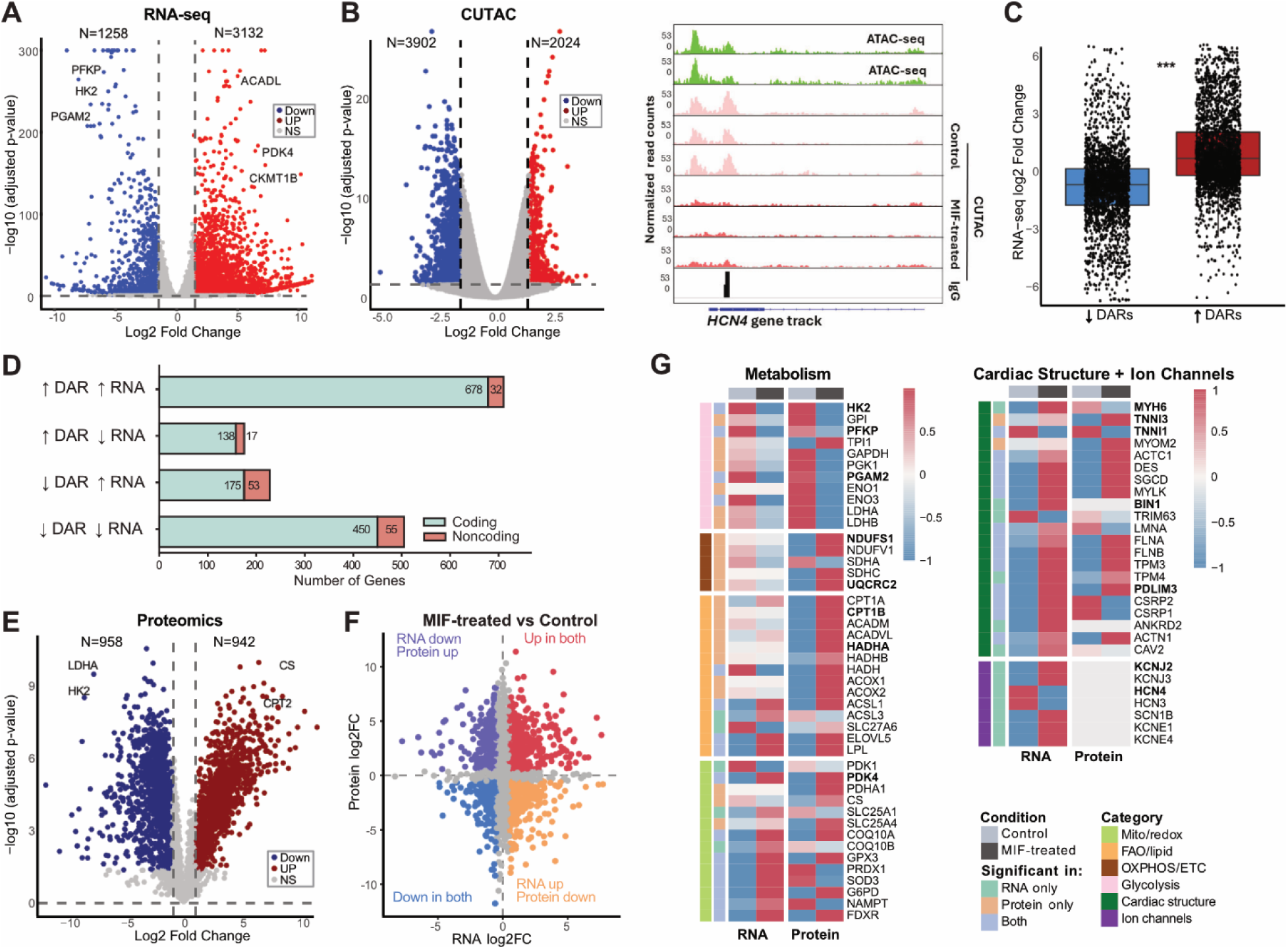
Multi-omics characterization of MIF-induced cardiomyocyte maturation reveals coordinated transcriptional, epigenetic, and proteomic remodeling. **(A)** RNA-seq volcano plot showing differentially expressed genes between MIF-treated and control hiPSC-CMs. **(B)** CUTAC volcano plot showing differentially accessible regions (DARs) between MIF-treated and control hiPSC-CMs. **(C)** Representative genome browser tracks at the HCN4 locus, with ATAC-seq tracks^40^ shown as a positive control for CUTAC in hiPSC-CMs. **(D)** Comparison of gene expression and chromatin accessibility (Wilcoxon rank-sum test, **p < 0.001). Boxplots show the median (center line), interquartile range (box limits), and whiskers extending to 1.5 × the interquartile range. Individual genes are shown as points. **(E)** Overlap between DAR-associated genes and differentially expressed genes categorized as coding and non-coding transcripts. **(F)** Proteomics volcano plot showing differentially expressed proteins between MIF-treated and control hiPSC-CMs. **(G)** Scatter plot comparing RNA and protein log2 fold changes in MIF-treated versus control hiPSC-CMs, highlighting concordant and discordant transcriptomic and proteomic changes. **(H)** Heatmaps showing selected metabolism-, cardiac structure-, and ion channel-related genes across RNA-seq and proteomics datasets. Statistical significance was defined as adjusted p ≤ 0.05 unless otherwise indicated (*n* = 3-4 independent differentiations).

In parallel, MIF-treated hiPSC-CMs exhibited altered expression of transcriptional regulators and chromatin-associated factors linked to metabolic gene control, including increased expression of PPARG, NR2F2, KLF9, and HOPX, alongside modulation of epigenetic regulators such as SMARCA2, TET1/2, and KAT2A (**Figure S2B**). Together, these transcriptional changes indicate that metabolic maturation is accompanied by broad remodeling of gene regulatory programs rather than isolated pathway activation.

To assess whether these transcriptional changes were supported by epigenetic remodeling, we profiled chromatin accessibility using CUTAC (three biological replicates per condition). Replicates clustered tightly, with MIF-treated samples forming a distinct group from controls (**Figure S2C**), with genome browser tracks showing concordance with previously reported ATAC-seq profiles in hiPSC-CMs^42^ (**Figure 2B**). Genome-wide analysis revealed a redistribution of accessible chromatin following MIF treatment, characterized by reduced accessibility at transcription start sites and increased accessibility at distal regulatory regions (**Figure S2D-E**). Differential accessibility analysis identified 5,926 regions with significant changes, including 3,902 regions with decreased and 2,024 regions with increased accessibility (**Figure 2B**).

Regions gaining accessibility were frequently associated with metabolic genes, including PDK4, mirroring transcriptional upregulation observed by RNA-seq (**Figure S2F**). In contrast, loci associated with immature cardiomyocyte programs, including the pacemaker gene HCN4, exhibited reduced accessibility (**Figure 2B**), consistent with transcriptional repression and a shift away from fetal-like metabolic and electrophysiological states. Globally, changes in chromatin accessibility correlated with transcriptional changes (**Figure 2C-D**), although discordant patterns were also observed, including enrichment for non-coding RNA loci, suggesting additional layers of post-transcriptional regulation.

We next assessed whether these changes were reflected at the protein level using whole-cell quantitative proteomics (four biological replicates per condition). PCA revealed strong clustering of replicates, with major variance in the treatment conditions (**Figure S2G**). MIF treatment resulted in significant changes in protein abundance, with 942 proteins upregulated and 958 downregulated relative to Day 30 control (**Figure 2E**). Consistent with transcriptomic and chromatin profiles, proteins involved in oxidative phosphorylation (CS, NDUFS1, UQCRC2) and fatty acid oxidation (HADHA, CPT1B) were increased, whereas glycolytic enzymes (LDHA, HK2) were reduced.

Integration of the three datasets revealed both concordant and discordant patterns across the different platforms (**Figure S2H-I**). A subset of genes showed consistent increases (n=181) or decreases (n=103) in transcript abundance and protein expression (**Figure 2F**), with concordantly upregulated genes enriched for mitochondrial and lipid metabolic processes and cardiac structural programs, and concordantly downregulated genes associated with glycolysis and biosynthetic processes. Notably, structural protein isoforms shifted toward a more mature profile, including increased TNNI3 and MYOM2 alongside reduced TNNI1 and MYOM1 expression (**Figure 2G**). Discordance between RNA and protein levels was also evident, consistent with post-transcriptional regulation during CM maturation. Across all layers, the dominant signal was a concerted remodeling toward suppression of hypoxia pathway, BMP signalling, and cell cycle while activating pathways involved in ECM organization, plasma membrane tubulation, and contraction (**Figure S2J-K**).

Collectively, these multi-omics analyses demonstrate that MIF-induced metabolic maturation is accompanied by global transcriptional, epigenetic, and proteomic reprogramming, reinforcing metabolic remodeling as a central and defining feature of this maturation paradigm.

### TNNT2-linked HCM variants exhibit impaired metabolic maturation

To determine how TNNT2-linked hypertrophic cardiomyopathy (HCM) variants affect CM metabolic maturation, we used genome-edited hiPSC lines harboring the I79N^+/−^ and R278C^+/−^ variants (hereafter I79N and R278C), together with their isogenic wild-type (WT) controls ^25,37^. All lines were differentiated into CMs and subjected to a four-week metabolic maturation (MM) regimen using the combined MIF medium described above, allowing comparison of immature (Day 30) and metabolic maturation (MM, Day 44) stages of CM development (**Figure 1A**).

We first assessed mitochondrial metabolic function using the Seahorse MitoStress assay in WT, R278C, and I79N hiPSC-CMs at Day 30 and after MM. As before, OCR and ECAR were measured (**Figure 3A**). At Day 30, both HCM variants exhibited significantly reduced mitochondrial respiration compared with WT hiPSC-CMs, as evidenced by decreased basal respiration, maximal respiration, spare respiratory capacity, and ATP production (**Figure 3B**). In parallel, the OCR/ECAR ratio was reduced in both variants (**Figure 3C**), indicating a relative shift away from oxidative metabolism toward glycolytic energy production. These findings demonstrate that TNNT2-linked HCM variants exhibit an early impairment in mitochondrial metabolic capacity during CM development.

**Figure 3.**
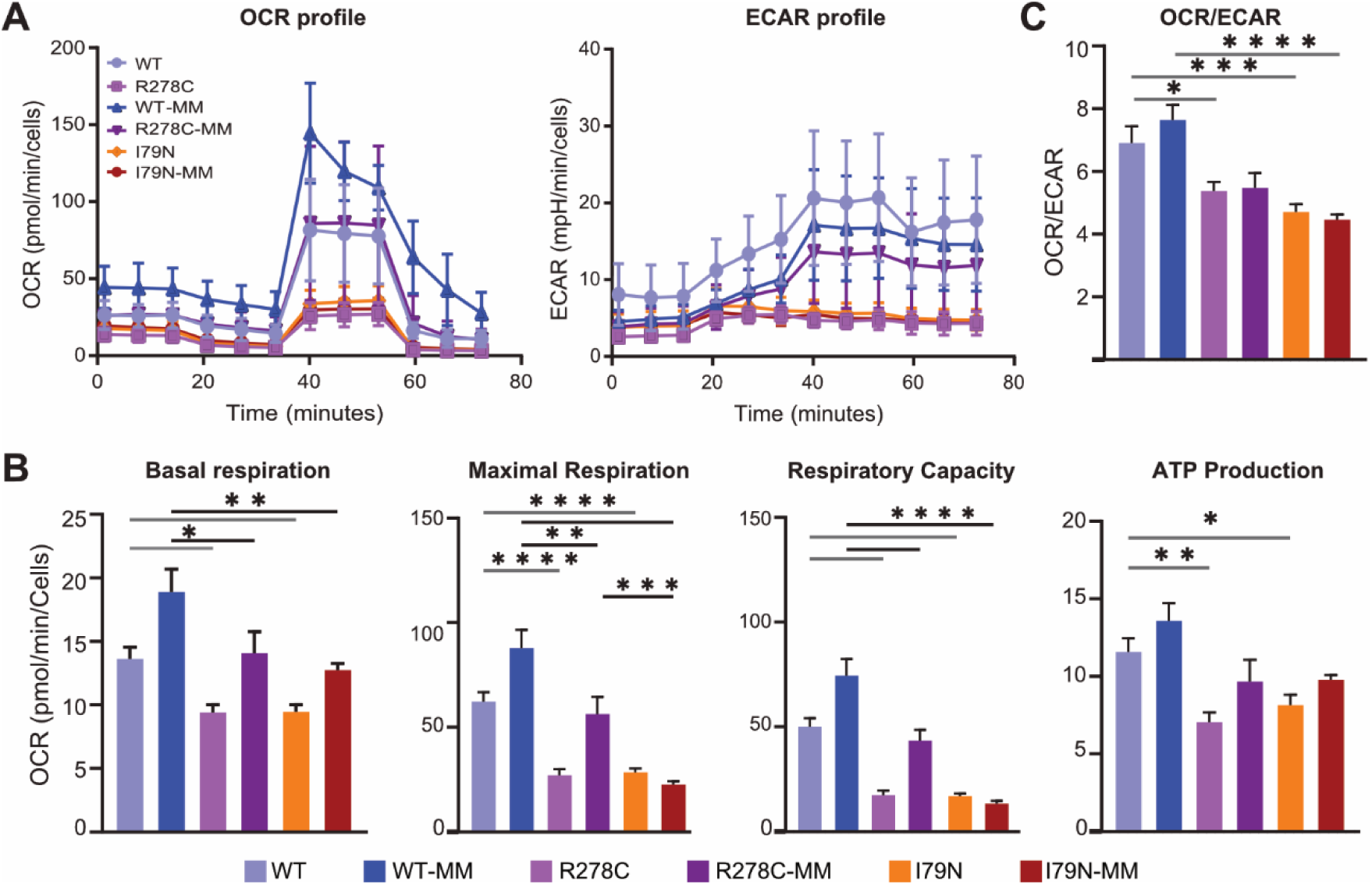
Metabolic characterization of TNNT2-linked HCM hiPSC-CMs during metabolic maturation using Seahorse Mito Stress Test analysis. **(A)** Oxygen consumption rate (OCR) and extracellular acidification rate (ECAR) profiles measured in WT, R278C, and I79N hiPSC-CMs under immature and metabolically matured (MM) conditions. **(B)** Quantification of basal respiration, maximal respiration, respiratory capacity, and ATP production derived from Seahorse mitochondrial stress tests. **(C)** OCR/ECAR ratio across experimental groups. Statistical significance was determined using one-way ANOVA with Tukey’s post-hoc test. Data are presented as mean ± SEM (*n* = 7-9 independent differentiations). \**p* ≤ 0.05, \*\***p** ≤ 0.01, \*\*\**p* ≤ 0.001, \*\*\*\***p** ≤ 0.0001.

Following metabolic maturation, WT hiPSC-CMs displayed the expected increase in mitochondrial respiratory parameters across all measured indices (**Figure 3B**). In contrast, both HCM variants remained metabolically compromised relative to WT MM-treated cells. The R278C-MM variant showed a partial improvement in mitochondrial respiration, consistent with a milder metabolic impairment, whereas the I79N-MM variant failed to substantially increase respiratory capacity following MM (**Figure 3B**). As a result, the metabolic profile of I79N-MM hiPSC-CMs remained similar to that observed at earlier developmental stages, indicating a persistent defect in metabolic maturation.

Together, these results demonstrate that TNNT2-linked HCM variants exhibit variant-specific defects in metabolic maturation, with the more clinically severe I79N variant displaying a pronounced and sustained impairment in mitochondrial oxidative metabolism. These findings support a model in which disrupted metabolic remodeling during CM maturation is a feature of TNNT2-linked HCM and scales with variant severity.

### Variant-specific transcriptional and proteomic remodeling emerges during metabolic maturation

We next examined how TNNT2 variants reshape gene regulatory programs across developmental stages by profiling transcriptomes at the immature (Day 30) and metabolically matured (Day 44, MM) states. PCA and pairwise correlation revealed that the largest source of variance across samples was metabolic maturation, followed by clear genotype-dependent separation. Notably, I79N samples consistently clustered furthest from WT and R278C (PC1 = 56%, PC2 = 28%; **Figure 4A**), indicating greater transcriptional divergence in the clinically more severe variant.

**Figure 4.**
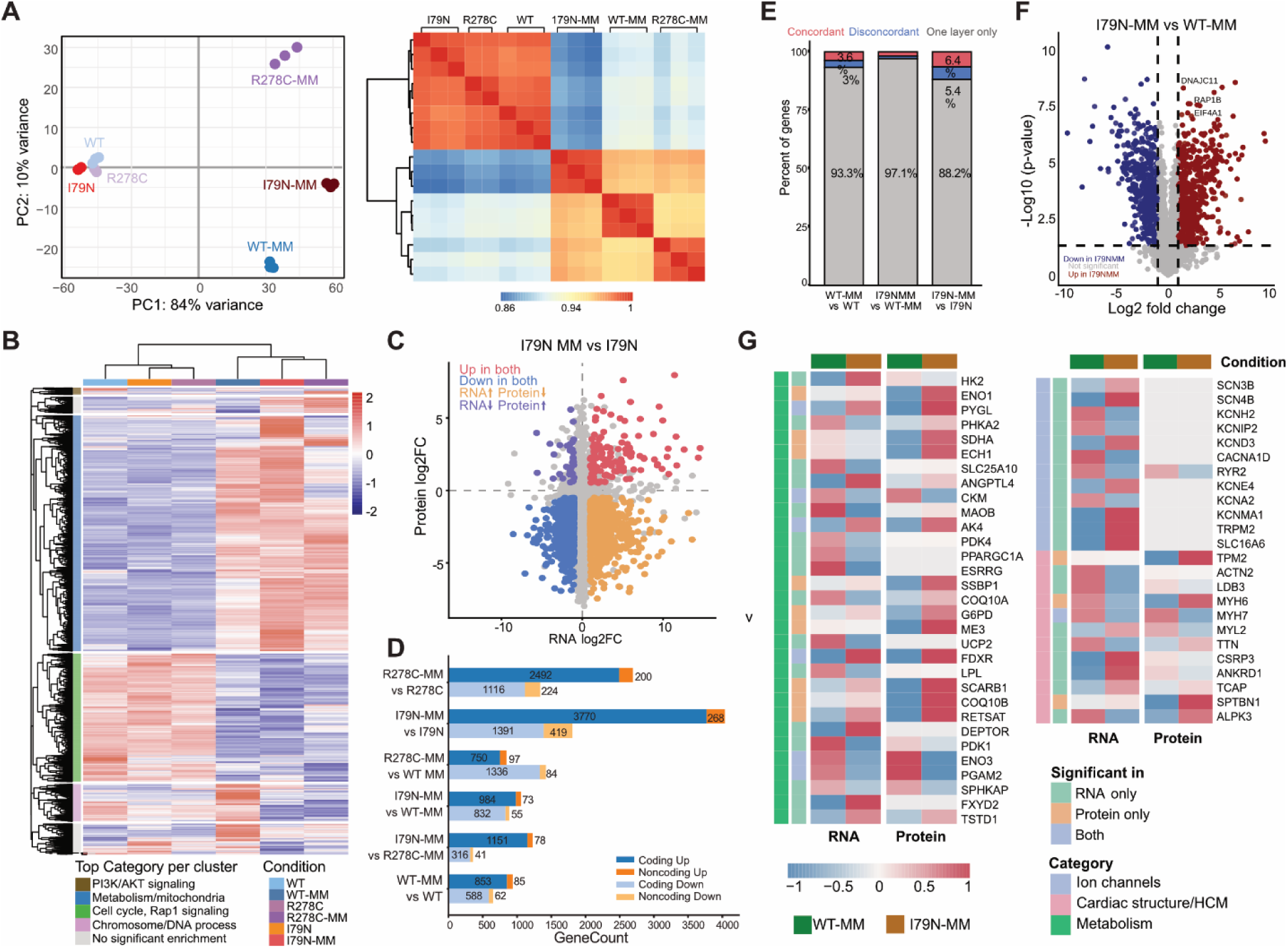
Transcriptomic and proteomic profiling of *TNNT2*-linked HCM hiPSC-CMs during metabolic maturation. **(A)** Principal component analysis (PCA) and sample correlation heatmap of RNA-seq datasets across WT, I79N, and R278C hiPSC-CMs under immature and metabolically matured (MM) conditions. **(B)** Heatmap of differentially expressed genes across experimental groups. **(C)** Scatter plot comparing RNA and protein log2 fold changes in I79N-MM versus I79N hiPSC-CMs, highlighting concordant and discordant transcriptomic and proteomic changes. **(D)** Numbers of coding and non-coding transcripts significantly up- or downregulated across indicated comparisons. **(E)** Overlap between RNA-seq and proteomics datasets showing concordant, discordant, and single-layer changes. **(F)** Proteomics volcano plot showing differentially expressed genes between I79N-MM and WT-MM hiPSC-CMs. **(G)** Heatmaps showing selected metabolism-, cardiac structure/HCM-, and ion channel-related genes across RNA-seq and proteomics datasets. Statistical significance: adjusted *p* ≤ 0.05 unless otherwise indicated (*n* = 3-4 independent differentiations for RNA-seq and proteomics).

To define global transcriptional trends, we compiled differentially expressed genes (DEGs) across all pairwise comparisons (∼8,000 genes) and performed hierarchical clustering across all conditions. This analysis revealed that metabolic maturation is the dominant driver of transcriptional remodeling irrespective of genotype, characterized by increased expression of metabolic genes and repression of cell cycle-associated genes (**Figure 4B**). These global trends mirror those observed in WT cells, indicating that core features of metabolic maturation are preserved in TNNT2 variants.

A second pattern emerged from clustering, where both TNNT2 variants grouped more closely with each other than with WT at both developmental stages, indicating a shared disease-associated transcriptional signature. At the immature stage (Day 30), however, transcriptional differences relative to WT were modest. Pairwise comparisons identified 141 upregulated and 423 downregulated genes in I79N, and 141 upregulated and 250 downregulated genes in R278C (**Figure S3A-B**). Direct comparison between variants revealed that I79N preferentially upregulated stress-associated pathways, including mitochondrial stress responses (**Figure S3C-D**), suggesting early activation of stress programs in the more severe variant.

In contrast, transcriptional differences became substantially more pronounced following metabolic maturation. Both variants exhibited extensive differential gene expression relative to WT-MM (I79N: 1,057 up and 887 down; R278C: 847 up and 1,420 down; **Figure S3E-F**). R278C-MM preferentially activated structural and contractile gene programs, whereas I79N-MM showed stronger activation of stress-responsive and extracellular matrix-associated pathways. Direct comparison of I79N-MM and R278C-MM further revealed suppression of sarcomere and contraction-related genes in I79N-MM alongside enrichment of adhesion and cytoskeletal remodeling programs, indicating divergent responses to metabolic and mechanical stress at later stages of maturation (**Figure S3G-H**).

To determine whether these transcriptional differences extend to the proteome, we performed quantitative proteomics on I79N hiPSC-CMs at Day 30 and after MM. Metabolic maturation induced widespread proteomic remodeling in I79N cells (265 upregulated and 3,756 downregulated proteins; **Figure S3I**). Compared to WT, MM treatment in I79N resulted in a disproportionate reduction in protein abundance, suggesting dysregulated proteome remodeling during maturation in the severe variant.

Comparison of RNA and protein changes revealed both concordant and discordant patterns (**Figure 4C**). While a subset of genes showed coordinated regulation, a substantial fraction exhibited increased RNA but decreased protein levels in I79N, indicating extensive post-transcriptional regulation. Consistent with this, non-coding RNA expression changes were more pronounced in I79N following MM (268 upregulated and 419 downregulated; **Figure 4D**), with distinct clustering patterns across samples (**Figure S3J**). Notably, concordance between RNA and protein changes was higher in within-sample comparisons (Day 30 vs MM) than in cross-genotype comparisons (I79N-MM vs WT-MM) (**Figure 4E**), suggesting that increased gene regulatory complexity is primarily associated with the maturation process rather than disease state alone.

We next assessed variant-specific proteomic changes following maturation by comparing I79N-MM to WT-MM (431 upregulated and 349 downregulated proteins; **Figure 4F**). I79N-MM cells exhibited reduced abundance of proteins involved in antioxidant defense and calcium handling, alongside increased expression of proteins linked to mitochondrial stress responses, translation, and signaling. These changes were accompanied by alterations in proteins related to metabolism, ion channels, and cardiac structure (**Figure 4G**), consistent with HCM-associated molecular remodeling.

Together, these findings demonstrate that metabolic maturation exposes and amplifies variant-specific regulatory programs in TNNT2-linked HCM. While early-stage differences are relatively modest, maturation reveals pronounced and multilayered transcriptional and proteomic remodeling, particularly in the I79N variant, highlighting the importance of developmental context in uncovering disease-relevant mechanisms.

### Chromatin remodeling partially explains transcriptional changes during metabolic maturation in HCM variants

To determine whether the observed transcriptional changes are accompanied by epigenetic remodeling, we profiled chromatin accessibility using CUTAC at Day 30 and after metabolic maturation (Day 44). Consistent with RNA-seq analyses, PCA and pair-wise correlation revealed only minor differences between WT and HCM variants at Day 30. In contrast, metabolic maturation emerged as the dominant source of variance, with additional genotype-dependent separation becoming more apparent after MM treatment (**Figure 5A, Figure S4A**). These patterns indicate that chromatin accessibility broadly tracks with transcriptional state, while also revealing variant-specific divergence upon maturation.

**Figure 5.**
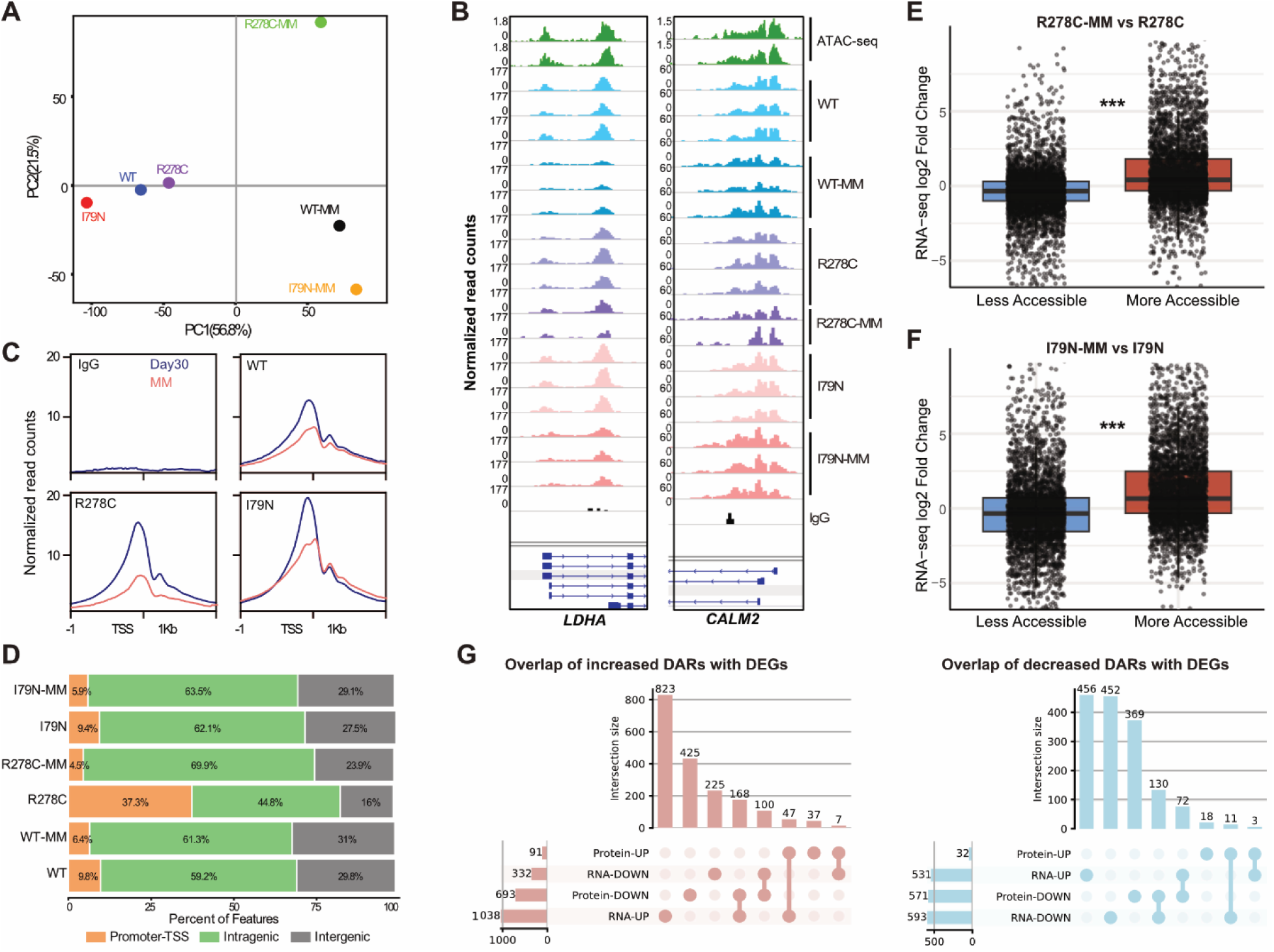
Chromatin accessibility remodeling during metabolic maturation of *TNNT2*-linked HCM hiPSC-CMs. **(A)** Principal component analysis (PCA) of CUTAC datasets across WT, I79N, and R278C hiPSC-CMs under immature and metabolically matured (MM) conditions. **(B)** Representative genome browser tracks showing chromatin accessibility at the *LDHA* and *CALM2* loci. ATAC-seq tracks^40^ are shown as positive controls for CUTAC. **(C)** CUTAC signal enrichment around transcription start sites (TSSs) in WT, I79N, and R278C hiPSC-CMs before and after metabolic maturation. IgG was used as a negative control. **(D)** Genomic annotation of CUTAC peaks showing promoter-TSS, intragenic, and intergenic distributions across experimental groups. **(E–F)** Comparison of gene expression and chromatin accessibility in R278C-MM vs R278C (E) and I79N-MM vs I79N (F) (Wilcoxon rank-sum test, \*\**p* < 0.001). Boxplots show the median (center line), interquartile range (box limits), and whiskers extending to 1.5 × the interquartile range. Individual genes are shown as points. **(G)** Overlap between DAR-associated genes and differentially expressed genes (DEGs) at the RNA and protein levels for regions with increased or decreased accessibility. Statistical significance: \**p* ≤ 0.05, \*\***p** ≤ 0.01, \*\*\****p*** ≤ 0.001, \*\*\*\***p** ≤ 0.0001 (*n* = 3 independent differentiations).

Inspection of CUTAC profiles at representative loci supported these global trends. Genome browser tracks at *CALM2*, involved in excitation-contraction coupling, and *LDHA*, a glycolysis-associated gene, showed strong concordance with previously published ATAC-seq datasets^40^ in WT hiPSC-CMs and revealed a general decrease in promoter-proximal accessibility following metabolic maturation (**Figure 5B**). To systematically assess this effect, we quantified the CUTAC signal across a 2 kb window centered on transcription start sites (TSSs) genome-wide. Both WT and HCM variants exhibited reduced TSS accessibility after MM treatment (**Figure 5C**), accompanied by a relative increase in accessibility within intronic and intergenic regions (**Figure 5D**), consistent with a global redistribution of accessible chromatin during maturation.

We next examined differential accessibility between variants and WT at the immature stage. At Day 30, I79N hiPSC-CMs exhibited a limited number of differentially accessible regions (DARs) relative to WT (n = 44), predominantly reflecting decreased accessibility (**Figure S4B**). In contrast, R278C hiPSC-CMs displayed a greater number of DARs (n = 115), with a bias toward increased accessibility (**Figure S4C**). These modest differences at the chromatin level mirror the relatively subtle transcriptional divergence observed at this stage.

In contrast, metabolic maturation induced extensive chromatin remodeling within each variant. Comparison of Day 44 MM samples to their Day 30 counterparts identified 5,956 increased and 5,849 decreased DARs in R278C, and 5,451 increased and 5,720 decreased DARs in I79N (**Figure S4D-G**). Changes in chromatin accessibility were positively correlated with changes in gene expression for both variants (**Figure 5E-F**), indicating that maturation-associated transcriptional remodeling is, in part, supported by epigenetic reprogramming.

To further resolve cross-layer relationships, we integrated chromatin accessibility, RNA-seq, and proteomics data in the I79N variant following MM treatment. UpSet analysis revealed substantial overlap between increased DARs and upregulated genes at the RNA (n = 1038) and protein (n = 91) levels individually, but more limited concordance across all three layers simultaneously (n = 47; **Figure 5G**). Concordance was observed to a greater extent for regions with decreased accessibility (n=130) (**Figure 5G**). Notably, a large fraction of genes exhibited discordant relationships between chromatin, RNA, and protein levels, consistent with an increased contribution of post-transcriptional regulation during later stages of maturation.

We next assessed variant-specific chromatin remodeling relative to WT after metabolic maturation. Both HCM variants displayed distinct accessibility profiles at Day 44, with I79N-MM showing 130 increased and 898 decreased DARs, and R278C-MM showing 364 increased and 158 decreased DARs relative to WT-MM (**Figure S4H-I**). Functional enrichment analysis of these DARs highlighted pathways associated with cardiomyopathy, including muscle filament sliding, sarcomere organization, and cardiac development, with stronger enrichment observed in I79N-MM (**Figure S4J-K**), consistent with its more severe phenotype.

Finally, integration of chromatin accessibility and RNA-seq data in HCM-MM versus WT-MM revealed limited global concordance between accessibility and transcriptional changes, in contrast to the stronger coupling observed during WT maturation. Nevertheless, a subset of disease-relevant genes, including regulators of contractile function and metabolism, exhibited coordinated changes across chromatin and transcriptional layers (**Figure S4L)**. Together, these findings indicate that while chromatin remodeling contributes to transcriptional reprogramming during metabolic maturation, its role is reduced in HCM, where additional regulatory mechanisms increasingly shape gene expression outputs.

### mTORC1 inhibition modulates maturation and disease-associated protein programs in I79N hiPSC-CMs

Previous studies have established that mTORC1 signaling maintains hiPSC-CMs in an immature, proliferative state, while its inhibition promotes CM maturation. In HCM samples, mTORC1 signaling is reported to be hyperactivated, contributing to pathological growth and remodeling ^30,33,43^. Using PROGENy pathway analysis^44^ of our RNA-seq dataset, we observed decreased expression of PI3K signaling genes in WT-MM cells relative to immature samples, suggesting altered activity of the PI3K-AKT-mTOR axis during metabolic maturation (**Figure S5A**). To determine how mTORC1 activity relates to cardiomyocyte maturation and TNNT2-linked HCM, we directly quantified mTORC1 signaling in our system.

We first utilized a validated bioluminescence resonance energy transfer (BRET) biosensor that reports mTORC1-mediated phosphorylation of a ULK1-derived peptide (T757)^45^ (**Figure 6A**). In WT hiPSC-CMs, treatment with the general mTOR inhibitor Torin1 significantly reduced the BRET signal, whereas a non-phosphorylatable (A757) biosensor control showed no response, confirming assay specificity (**Figure 6B**).

**Figure 6.**
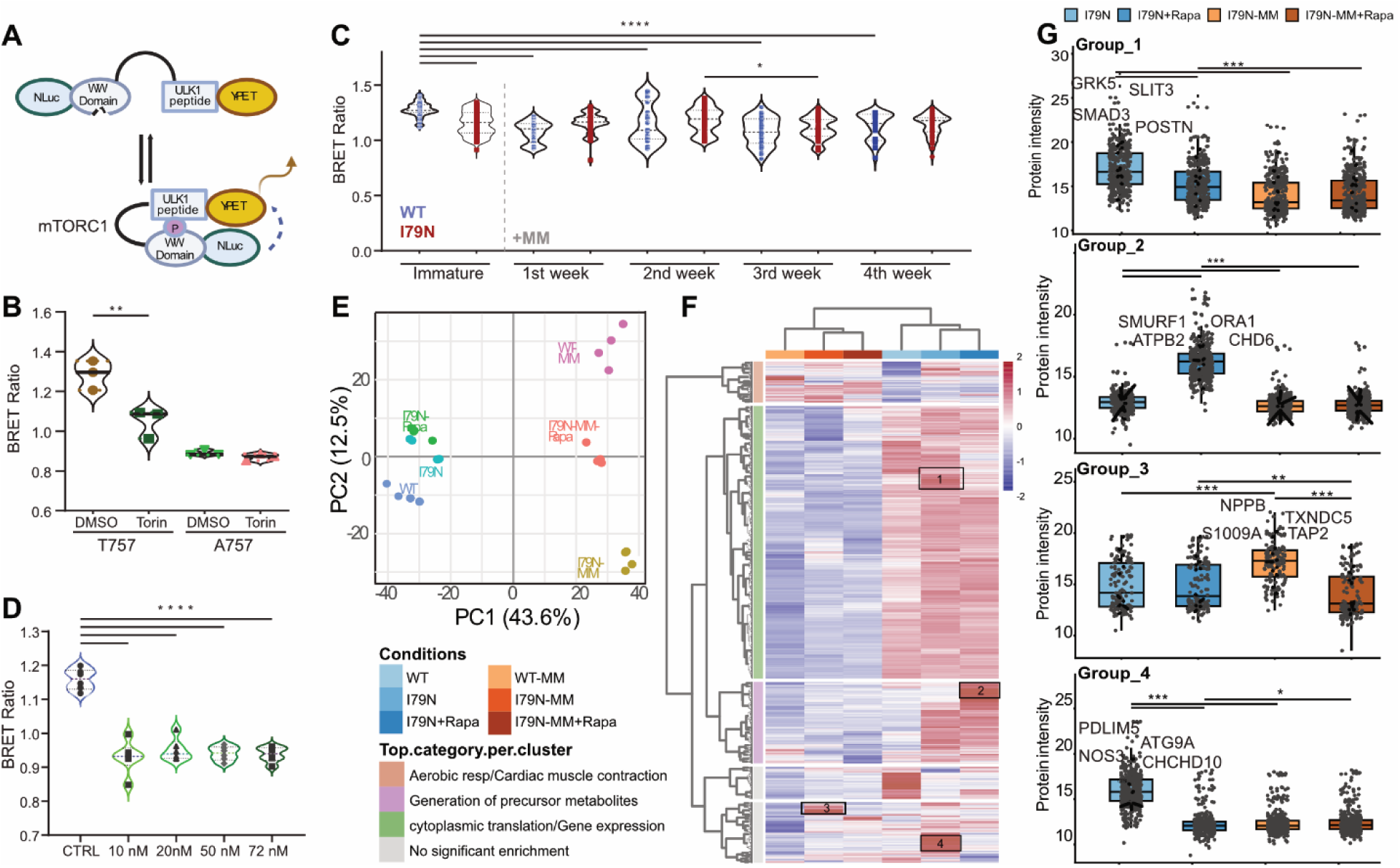
Effects of mTORC1 inhibition on *TNNT2*-linked HCM hiPSC-CMs during metabolic maturation. **(A)** Schematic of the mTORC1 biosensor used to monitor mTORC1-ULK1 signaling activity through BRET. **(B)** Validation of the biosensor using T757 and A757 control constructs following Torin treatment (*n*=3 independent differentiation). **(C)** Longitudinal measurement of mTORC1 activity during metabolic maturation in WT and I79N hiPSC-CMs (*n*=5 independent differentiations). **(D)** Dose-response analysis of rapamycin (Rapa) treatment using the mTORC1 biosensor (*n* = 3 independent differentiations). Statistical significance for B-D was determined using one-way ANOVA with Tukey’s post-hoc test. Data are presented as mean ± SEM. \**p* ≤ 0.05, \*\***p** ≤ 0.01, \*\*\****p*** ≤ 0.001, \*\*\*\***p** ≤ 0.0001. **(E)** Principal component analysis (PCA) of proteomic datasets across WT, I79N, I79N+Rapa, WT-MM, I79N-MM, and I79N-MM+Rapa hiPSC-CMs. **(F)** Heatmap of differentially expressed proteins across experimental groups. **(G)** Protein expression changes within selected clusters identified by hierarchical clustering (*n*= 4 independent differentiations for proteomics).

Monitoring mTORC1 dynamics during maturation revealed that WT activity decreased sharply within the first week of MM treatment and remained suppressed throughout (**Figure 6C**), identifying mTORC1 downregulation as an early, sustained feature of metabolic maturation. Interestingly, I79N hiPSC-CMs exhibited reduced mTORC1 activity relative to WT at the immature stage, which remained low post-maturation (**Figure 6C**). This baseline suppression likely reflects the impaired mitochondrial function in I79N cells (**Figure 3B)**. The resulting energetic stress may contribute to AMPK-dependent inhibition of mTORC1, as supported by increased expression of the AMP sensor AK4 and the mTORC1 inhibitor DDIT4 (**Figure S5B**),), which has been linked to biosynthetic plasticity during the stress response ^46–48^. This suggests that while advanced HCM is characterized by mTORC1 hyperactivation, the early stages of the I79N variant may be characterized by suppression of growth signaling. While direct measurement of the other mTORC1 targets such as S6K phosphorylation was beyond the scope of this study, the observed upregulation of S6K protein alongside significant downregulation of IRS1 at the protein level in I79N cells may reflect activation of an S6K to IRS1 negative feedback loop, potentially promoting anabolic resistance ^49^. Rapamycin-mediated rescue of IRS1 protein levels further supports this interpretation (**Figure S5C**).

To test the functional impact of this signaling axis, we identified 10 nM as the minimal effective concentration of rapamycin for sustained mTORC1 suppression (**Figure 6D**). We then assessed how mTORC1 inhibition reshapes the proteomic landscape in I79N hiPSC-CMs by performing quantitative proteomics in four biological replicates following rapamycin treatment during two distinct windows: an early intervention (Day 16-30) and a late intervention during metabolic maturation (Day 30-44). PCA of these four samples along with WT and WT MM revealed clear separation of treatment conditions (**Figure 6E**). Although the major degree of variance is due to MM treatment, WT MM, I79N MM and I79N MM + Rapamycin showed distinct clustering along PC2 axis, indicating robust proteome remodeling in response to mTORC1 inhibition.

Early rapamycin treatment altered 634 proteins, characterized by downregulation of fetal sarcomere components (e.g., MYH3) and regulators of lipid and nucleotide metabolism, alongside upregulation of proteins involved in PI3K/AKT pathway modulation, lipid metabolism, and protein quality control (**Figure S5D**), suggesting that intervention at the immature stage affected maturation and metabolism pathways. Late rapamycin treatment resulted in 682 differentially expressed proteins, including reduced abundance of mitochondrial and cholesterol biosynthesis proteins and increased expression of mature sarcomeric components such as MYOM2. In addition, isoform shifts were observed in proteins involved in vesicular trafficking (RAB1A to RAB1B) and PKA signaling (PRKAR2A to PRKAR2B) (**Figure S5E-G**), suggesting remodeling of intracellular signaling networks relevant to cardiomyocyte function.

Hierarchical clustering across all conditions identified four distinct protein groups regulated by genotype and rapamycin treatment (**Figure 6H**). Groups 1 and 4 exhibited elevated expression in I79N hiPSC-CMs compared to WT controls, which were markedly reduced following Rapamycin treatment. They included markers of hypertrophic stress (*GRK5, SMAD3*), pro-fibrotic signaling (*POSTN, SLIT3*), and proteostasis imbalance (*CHCHD10, ATG9A*) (**Figure 6G**). Conversely, Group 2 proteins were expressed at low levels in both WT and I79N hiPSC-CMs under basal conditions but were robustly upregulated following Rapamycin treatment. This group was enriched for regulators of calcium handling (*ORAI1, ATP2B3*), transcriptional regulation (*CHD6*), and ER-associated degradation (*DERL2*) (**Figure 6G**). Finally, Group 3 proteins showed low expression in WT MM but were specifically induced in I79N MM cells before being suppressed by rapamycin. This group included pathological markers like BNP (*NPPB*) and oxidative stress mediators (*TXNDC5*) (**Figure 6G**). The selective induction of these proteins upon metabolic maturation in I79N cells suggests that increased energetic demand exacerbates an underlying mutation-driven stress program, which remains dependent on mTORC1 activity.

Together, these data indicate that mTORC1 inhibition performs a dual role: it suppresses a cardiomyocyte-intrinsic disease program, spanning hypertrophic signaling and structural remodeling, while simultaneously inducing an adaptive program that enhances calcium handling and protein quality control. These findings support a model in which mTORC1 integrates sarcomere mutation-induced stress with metabolic state to drive early remodeling in I79N cardiomyocytes, and that its inhibition shifts cells toward a more homeostatic and stress-resilient phenotype.

## Discussion

In this study, we present a comprehensive multi-omic map of metabolic maturation in hiPSC-CMs and reveal how this developmental transition serves as a critical checkpoint for the manifestation of TNNT2-linked HCM. By integrating transcriptomic, epigenetic, and proteomic profiling, we demonstrate that metabolic maturation is not merely a shift in substrate utilization, but a coordinated global reprogramming event. Our findings suggest that the metabolic state of the cardiomyocyte is a primary determinant of disease penetrance, where the transition to oxidative metabolism unmasks variant-specific defects that remain latent in immature cells.

While many studies focus on the structural maturation gap, our data show that the metabolic transition induced by MIF treatment triggers a profound, multi-layered reprogramming of the cellular landscape. The high degree of coordination between chromatin accessibility (CUTAC), mRNA, and protein abundance of metabolic and structural programs suggests that metabolic maturation is a hard-coded developmental program. The observed global reduction in TSS accessibility mirrors the transition to repressive chromatin states seen in embryonic heart development as cells acquire specialized identities ^50–52^. Furthermore, this finding suggests that metabolic maturation is associated with suppression of signaling pathways such as hypoxia and BMP signaling, the latter showing a stage-specific role in CM development ^53,54^. Persistent induction of hypoxia responses remains a major limitation in the current hiPSC-CM maturation field, and efforts to overcome this limitation have focused on the development of bioreactors and optimized medium depth in cell culture ^6,55^.

A central finding of this work is that TNNT2-linked variants, particularly the severe I79N mutation, exhibit relatively modest molecular divergence at the immature stage. Rather, the energetic challenge of metabolic maturation serves as the second hit that manifests more disease-related molecular patterns. Specifically, we observed an early decrease in mitochondrial respiration that scales with variant severity. The concordant repression of CKMT2 (mitochondrial creatine kinase) across all datasets lends strong support to the ATP depletion theory of HCM ^56,57^. The upregulation of ITLN1 and AK4 likely represents a protective sensing response via AMPK signaling as the cell attempts to mitigate this energetic stress^48,58,59^.

While R278C hiPSC-CMs showed a milder metabolic phenotype and a largely “shared” disease signature, the I79N variant activated distinct stress-response pathways, including early induction of hypoxia, lipid biosynthesis, and contractile programs. The activation of lipid biosynthesis pathways may represent an adaptive response to support membrane biogenesis and repair required to maintain cellular integrity under mechanical stress ^60,61^. This observation is consistent with previous studies of R92W-TNNT2-linked HCM in mouse models^62^. Consistently, lipid biosynthesis regulators such as ACLY were upregulated. As ACLY utilizes TCA cycle-derived citrate to generate cytosolic acetyl-CoA, enhanced lipid biosynthesis may promote TCA cycle cataplerosis, potentially contributing to the persistent mitochondrial respiration defect observed in I79N cells following maturation ^63^. In addition, ACLY-derived acetyl-CoA can serve as a substrate for histone acetylation, raising the possibility that early metabolic reprogramming contributes to the chromatin remodeling observed in mature HCM variants ^64^. Previous studies have shown that patients who carried a large effect size pathogenic HCM variant along with R278C^+/-^ had notably worse outcomes, including heart failure-related events, compared to carriers of large effect size pathogenic variants alone ^65,66^. Our finding supports the clinical observation that R278C may act as a phenotypic amplifier when combined with other variants, whereas I79N drives a severe, early-onset phenotype through independent transcriptional trajectories.

Our investigation into mTORC1 signaling reveals a striking departure from the hyperactivation typically associated with clinical HCM. We observed that mTORC1 activity is downregulated early in WT maturation and remains suppressed, a pattern that aligns with studies showing that mTOR inhibition is a requirement for functional maturity and cellular quiescence^30,43,67^. In the I79N variant, this suppression is even more pronounced, likely reflecting an early compensatory response to preserve ATP by redirecting energy away from biosynthetic growth programs and toward stress response pathways, as described in prior work on metabolic plasticity in stressed cells^47^. This finding is also consistent with the absence of overt ventricular hypertrophy in TNNT2-linked HCM, which stands in contrast to the hypertrophic remodeling more commonly observed in MYBPC3-linked HCM ^12,68^. Haploinsufficiency in MYBPC3-linked variants reduces sarcomeric protein dosage, necessitating compensatory upregulation of protein synthesis that would require mTORC1 activation ^69^. By contrast, the poison-peptide mechanism of *TNNT2* variants^70^ may render anabolic mTORC1 signaling dispensable at this stage. This variant-specific difference may explain why mTORC1 activity is suppressed rather than elevated in I79N cells, and why the ventricular hypertrophy is less pronounced in TNNT2-linked HCM patients^71,72^. However, the dysregulation of the MAPK/ERK cascade specifically during maturation in HCM cells^73,74^ indicates that the signaling logic of the cell is beginning to rewire toward pathological growth, even while growth-promoting mTORC1 remains suppressed.

The timing of rapamycin intervention provides a compelling argument for the plasticity of the HCM phenotype. Rapamycin treatment successfully reversed the fetal switch (e.g., *MYH3, MYOM2*), and decreased NPPB (BNP), a primary clinical biomarker for HCM severity^75^. Beyond suppressing disease markers, mTORC1 inhibition induced an adaptive program (Group 2) that enhanced calcium handling and proteostasis (via *DERL2* and *DNAJC5*). This result suggests that pharmacological targeting of mTORC1, or the downstream induction of autophagy ^76,77^, does not simply blunt hypertrophy but actively shifts the cell toward a more stress-resilient, adult-like homeostatic state.

By defining the molecular trajectories of metabolic maturation, we have identified a critical developmental window where genetic predisposition intersects with physiological demand. Our results demonstrate that TNNT2-linked HCM is a dynamic process that can be modulated by metabolic and signaling interventions. The ability to reverse maturation-exacerbated stress programs through temporal mTORC1 control provides a proof-of-concept for metabolic-based therapeutics, emphasizing the need for developmentally attuned models in the quest for precision cardiovascular medicine.

## Acknowledgement

Methodology: H.H.; Investigation: H.H. F.J. (Proteomics);S.S(Genome editing), Visualization: H.H.; Funding acquisition: S.S.T. and G.F.T.; Supervision: S.S.T. and G.F.T.; Writing-original draft: H.H.; Writing-review and editing: S.S.T., and G.F.T.

## Data availability

All sequencing data have been deposited in the Gene Expression Omnibus (GEO) under accession numbers GSE307649 and GSE307652 and are accessible to reviewers using the reviewer access token wbyhmkyyfpohpgf.

## References

1. Woodruff RC, Tong X, Loustalot FV, Khan SS, Shah NS, Jackson SL, Vaughan AS. Cardiovascular Disease Mortality Trends, 2010–2022: An Update with Final Data. American Journal of Preventive Medicine. 2025;68(2):391–395.

2. Yildirim Z, Swanson K, Wu X, Zou J, Wu J. Next-Gen Therapeutics: Pioneering Drug Discovery with iPSCs, Genomics, AI, and Clinical Trials in a Dish. Annual Review of Pharmacology and Toxicology. 2025;65(1):71–90.

3. Ahmed RE, Anzai T, Chanthra N, Uosaki H. A Brief Review of Current Maturation Methods for Human Induced Pluripotent Stem Cells-Derived Cardiomyocytes. Frontiers in Cell and Developmental Biology. 2020;8:178.

4. Li W, Luo X, Strano A, Arun S, Gamm O, Poetsch MS, Hasse M, Steiner R-P, Fischer K, Pöche J, Ulbricht Y, Lesche M, Trimaglio G, El-Armouche A, Dahl A, et al. Comprehensive promotion of iPSC-CM maturation by integrating metabolic medium with nanopatterning and electrostimulation. Nature Communications. 2025;16(1):2785.

5. Ronaldson-Bouchard K, Ma SP, Yeager K, Chen T, Song L, Sirabella D, Morikawa K, Teles D, Yazawa M, Vunjak-Novakovic G. Advanced maturation of human cardiac tissue grown from pluripotent stem cells. Nature. 2018;556(7700):239–243.

6. Prondzynski M, Berkson P, Trembley MA, Tharani Y, Shani K, Bortolin RH, Sweat ME, Mayourian J, Yucel D, Cordoves AM, Gabbin B, Hou C, Anyanwu NJ, Nawar F, Cotton J, et al. Efficient and reproducible generation of human iPSC-derived cardiomyocytes and cardiac organoids in stirred suspension systems. Nature Communications. 2024;15(1):5929.

7. Maaref Y, Jannati S, Jayousi F, Lange P, Akbari M, Chiao M, Tibbits GF. Developing a soft micropatterned substrate to enhance maturation of human induced pluripotent stem cell-derived cardiomyocytes. Acta Biomaterialia. 2024;190:133–151.

8. Pocock MW, Reid JD, Robinson HR, Charitakis N, Krycer JR, Foster SR, Fitzsimmons RL, Lor M, Devilée LAC, Batho CAP, Tuano N, Howden SE, Vlahos K, Watt KI, Piers AT, et al. Maturation of human cardiac organoids enables complex disease modeling and drug discovery. Nature Cardiovascular Research. 2025;4(7):821–840.

9. Feyen DAM, McKeithan WL, Bruyneel AAN, Spiering S, Hörmann L, Ulmer B, Zhang H, Briganti F, Schweizer M, Hegyi B, Liao Z, Pölönen R-P, Ginsburg KS, Lam CK, Serrano R, et al. Metabolic Maturation Media Improve Physiological Function of Human iPSC-Derived Cardiomyocytes. Cell Reports. 2020;32(3):107925.

10. Wang L, Wada Y, Ballan N, Schmeckpeper J, Huang J, Rau CD, Wang Y, Gepstein L, Knollmann BC. Triiodothyronine and dexamethasone alter potassium channel expression and promote electrophysiological maturation of human-induced pluripotent stem cell-derived cardiomyocytes. Journal of Molecular and Cellular Cardiology. 2021;161:130–138.

11. Maas RGC, Van Den Dolder FW, Yuan Q, Van Der Velden J, Wu SM, Sluijter JPG, Buikema JW. Harnessing developmental cues for cardiomyocyte production. Development. 2023;150(15):dev201483.

12. Birket MJ, Ribeiro MC, Kosmidis G, Ward D, Leitoguinho AR, van de Pol V, Dambrot C, Devalla HD, Davis RP, Mastroberardino PG, Atsma DE, Passier R, Mummery CL. Contractile Defect Caused by Mutation in MYBPC3 Revealed under Conditions Optimized for Human PSC-Cardiomyocyte Function. Cell Reports. 2015;13(4):733–745.

13. Miki K, Deguchi K, Nakanishi-Koakutsu M, Lucena-Cacace A, Kondo S, Fujiwara Y, Hatani T, Sasaki M, Naka Y, Okubo C, Narita M, Takei I, Napier SC, Sugo T, Imaichi S, et al. ERRγ enhances cardiac maturation with T-tubule formation in human iPSC-derived cardiomyocytes. Nature Communications. 2021;12(1):3596.

14. Fu K, Nakano H, Morselli M, Chen T, Pappoe H, Nakano A, Pellegrini M. A temporal transcriptome and methylome in human embryonic stem cell-derived cardiomyocytes identifies novel regulators of early cardiac development. Epigenetics. 2018;13(10–11):1013–1026.

15. Krup AL, Winchester SAB, Ranade SS, Agrawal A, Devine WP, Sinha T, Choudhary K, Dominguez MH, Thomas R, Black BL, Srivastava D, Bruneau BG. A *Mesp1*-dependent developmental breakpoint in transcriptional and epigenomic specification of early cardiac precursors. Development. 2023;150(9):dev201229.

16. Liu Q, Jiang C, Xu J, Zhao M-T, Van Bortle K, Cheng X, Wang G, Chang HY, Wu JC, Snyder MP. Genome-Wide Temporal Profiling of Transcriptome and Open Chromatin of Early Cardiomyocyte Differentiation Derived From hiPSCs and hESCs. Circulation Research. 2017;121(4):376–391.

17. Pereira IT, Spangenberg L, Robert AW, Amorín R, Stimamiglio MA, Naya H, Dallagiovanna B. Cardiomyogenic differentiation is fine-tuned by differential mRNA association with polysomes. BMC Genomics. 2019;20(1):219.

18. Uosaki H, Cahan P, Lee DI, Wang S, Miyamoto M, Fernandez L, Kass DA, Kwon C. Transcriptional Landscape of Cardiomyocyte Maturation. Cell Reports. 2015;13(8):1705–1716.

19. Pepin ME, Gong X, Schulze A, Backs J. Metabo-epigenetic circuits of heart failure: chromatin-modifying enzymes as determinants of metabolic plasticity. EMBO Molecular Medicine. 2025;18(2):362–380.

20. Li X, Egervari G, Wang Y, Berger SL, Lu Z. Regulation of chromatin and gene expression by metabolic enzymes and metabolites. Nature Reviews Molecular Cell Biology. 2018;19(9):563–578.

21. Chen L, Chen M, Yang X, Hu Y, Qiu C, Fu Y, Lan X, Luo G, Liu Q, Liu M. Energy metabolism in cardiovascular diseases: unlocking the hidden powerhouse of cardiac pathophysiology. Frontiers in Endocrinology. 2025;16:1617305.

22. Marian AJ, Braunwald E. Hypertrophic Cardiomyopathy: Genetics, Pathogenesis, Clinical Manifestations, Diagnosis, and Therapy. Circulation Research. 2017;121(7):749–770.

23. Wei B, Jin J-P. TNNT1, TNNT2, and TNNT3: Isoform genes, regulation, and structure–function relationships. Gene. 2016;582(1):1–13.

24. Knollmann BC, Kirchhof P, Sirenko SG, Degen H, Greene AE, Schober T, Mackow JC, Fabritz L, Potter JD, Morad M. Familial Hypertrophic Cardiomyopathy-Linked Mutant Troponin T Causes Stress-Induced Ventricular Tachycardia and Ca^2+^-Dependent Action Potential Remodeling. Circulation Research. 2003;92(4):428–436.

25. Shafaattalab S, Li AY, Gunawan MG, Kim B, Jayousi F, Maaref Y, Song Z, Weiss JN, Solaro RJ, Qu Z, Tibbits GF. Mechanisms of Arrhythmogenicity of Hypertrophic Cardiomyopathy-Associated Troponin T (TNNT2) Variant I79N. Frontiers in Cell and Developmental Biology. 2021;9:787581.

26. Wang L, Kim K, Parikh S, Cadar AG, Bersell KR, He H, Pinto JR, Kryshtal DO, Knollmann BC. Hypertrophic cardiomyopathy-linked mutation in troponin T causes myofibrillar disarray and pro-arrhythmic action potential changes in human iPSC cardiomyocytes. Journal of Molecular and Cellular Cardiology. 2018;114:320–327.

27. Dababneh S, Hamledari H, Maaref Y, Jayousi F, Hosseini DB, Khan A, Jannati S, Jabbari K, Arslanova A, Butt M, Roston TM, Sanatani S, Tibbits GF. Advances in Hypertrophic Cardiomyopathy Disease Modelling Using hiPSC-Derived Cardiomyocytes. Canadian Journal of Cardiology. 2024;40(5):766–776.

28. Nollet EE, Sequeira V, Maack C, Ochala J, Van Der Velden J. Metabolic alterations in human hypertrophic cardiomyopathy. The Journal of Cardiovascular Aging. 2025;5(2).

29. Laplante M, Sabatini DM. mTOR Signaling in Growth Control and Disease. Cell. 2012;149(2):274–293.

30. Garbern JC, Helman A, Sereda R, Sarikhani M, Ahmed A, Escalante GO, Ogurlu R, Kim SL, Zimmerman JF, Cho A, MacQueen L, Bezzerides VJ, Parker KK, Melton DA, Lee RT. Inhibition of mTOR Signaling Enhances Maturation of Cardiomyocytes Derived From Human-Induced Pluripotent Stem Cells via p53-Induced Quiescence. Circulation. 2020;141(4):285–300.

31. Gao X-M, Wong G, Wang B, Kiriazis H, Moore X-L, Su Y-D, Dart A, Du X-J. Inhibition of mTOR reduces chronic pressure-overload cardiac hypertrophy and fibrosis. Journal of Hypertension. 2006;24(8):1663–1670.

32. McMullen JR, Sherwood MC, Tarnavski O, Zhang L, Dorfman AL, Shioi T, Izumo S. Inhibition of mTOR Signaling With Rapamycin Regresses Established Cardiac Hypertrophy Induced by Pressure Overload. Circulation. 2004;109(24):3050–3055.

33. Davogustto GE, Salazar RL, Vasquez HG, Karlstaedt A, Dillon WP, Guthrie PH, Martin JR, Vitrac H, De La Guardia G, Vela D, Ribas-Latre A, Baumgartner C, Eckel-Mahan K, Taegtmeyer H. Metabolic remodeling precedes mTORC1-mediated cardiac hypertrophy. Journal of Molecular and Cellular Cardiology. 2021;158:115–127.

34. Sciarretta S, Forte M, Frati G, Sadoshima J. New Insights Into the Role of mTOR Signaling in the Cardiovascular System. Circulation Research. 2018;122(3):489–505.

35. Beltrami M, Fedele E, Fumagalli C, Mazzarotto F, Girolami F, Ferrantini C, Coppini R, Tofani L, Bertaccini B, Poggesi C, Olivotto I. Long-Term Prevalence of Systolic Dysfunction in MYBPC3 Versus MYH7-Related Hypertrophic Cardiomyopathy. Circulation: Genomic and Precision Medicine. 2023;16(4):363–371.

36. Bu H, Ding Y, Li J, Zhu P, Shih Y-H, Wang M, Zhang Y, Lin X, Xu X. Inhibition of mTOR or MAPK ameliorates vmhcl/myh7 cardiomyopathy in zebrafish. JCI Insight. 2021;6(24):e154215.

37. Shafaatalab S, Li AY, Jayousi F, Maaref Y, Dababneh S, Hamledari H, Baygi DH, Barszczewski T, Ruprai B, Jannati S, Nagalingam R, Cool AM, Langa P, Chiao M, Roston T, et al. Mechanisms of Pathogenicity of Hypertrophic Cardiomyopathy-Associated Troponin T (TNNT2) Variant R278C+/- During Development. 2023.

38. Henikoff S, Henikoff J, Ahmad K. Simplified Epigenome Profiling Using Antibody-tethered Tagmentation. BIO-PROTOCOL. 2021;11(11).

39. Gomez-Garcia MJ, Quesnel E, Al-attar R, Laskary AR, Laflamme MA. Maturation of human pluripotent stem cell derived cardiomyocytes in vitro and in vivo. Seminars in Cell & Developmental Biology. 2021;118:163–171.

40. Lian X, Zhang J, Azarin SM, Zhu K, Hazeltine LB, Bao X, Hsiao C, Kamp TJ, Palecek SP. Directed cardiomyocyte differentiation from human pluripotent stem cells by modulating Wnt/β-catenin signaling under fully defined conditions. Nature Protocols. 2013;8(1):162–175.

41. Cardoso AC, Lam NT, Savla JJ, Nakada Y, Pereira AHM, Elnwasany A, Menendez-Montes I, Ensley EL, Petric UB, Sharma G, Sherry AD, Malloy CR, Khemtong C, Kinter MT, Tan WLW, et al. Mitochondrial Substrate Utilization Regulates Cardiomyocyte Cell Cycle Progression. Nature Metabolism. 2020;2(2):167–178.

42. Bertero A, Fields PA, Ramani V, Bonora G, Yardimci GG, Reinecke H, Pabon L, Noble WS, Shendure J, Murry CE. Dynamics of genome reorganization during human cardiogenesis reveal an RBM20-dependent splicing factory. Nature Communications. 2019;10(1):1538.

43. Tian Y, Lucena-Cacace A, Tani K, Elvandari AP, Allendes Osorio RS, Narita M, Matsumura Y, Paixao IC, Miyoshi Y, Inagaki A, Junghof J, Yoshida Y. Generation of mature epicardium derived from human-induced pluripotent stem cells via inhibition of mTOR signaling. Nature Communications. 2025;16(1):5902.

44. Schubert M, Klinger B, Klünemann M, Sieber A, Uhlitz F, Sauer S, Garnett MJ, Blüthgen N, Saez-Rodriguez J. Perturbation-response genes reveal signaling footprints in cancer gene expression. Nature Communications. 2018;9(1):20.

45. Bouquier N, Moutin E, Tintignac LA, Reverbel A, Jublanc E, Sinnreich M, Chastagnier Y, Averous J, Fafournoux P, Verpelli C, Boeckers T, Carnac G, Perroy J, Ollendorff V. AIMTOR, a BRET biosensor for live imaging, reveals subcellular mTOR signaling and dysfunctions. BMC biology. 2020;18(1):81.

46. Foltyn M, Luger A-L, Lorenz NI, Sauer B, Mittelbronn M, Harter PN, Steinbach JP, Ronellenfitsch MW. The physiological mTOR complex 1 inhibitor DDIT4 mediates therapy resistance in glioblastoma. British Journal of Cancer. 2019;120(5):481–487.

47. Michalski C, Cheung C, Oh JH, Ackermann E, Popescu CR, Archambault A-S, Prusinkiewicz MA, Da Silva R, Majdoubi A, Viñeta Paramo M, Xu RY, Reicherz F, Patterson AE, Golding L, Sharma AA, et al. DDIT4L regulates mitochondrial and innate immune activities in early life. JCI Insight. 2024;9(5):e172312.

48. Zhang S-M, Luo H, Li Z, Du M, Chen J, Jiang D-S, Zhang Y, Li C, Yang X, Wang X-S, Fang Z-M, Gong F-H, Yang J. Adenylate kinase 4 (AK4) deficiency prevents vascular smooth muscle cell phenotypic switching by regulating mitochondrial dysfunction through AMPKα inactivation. Atherosclerosis. 2025:120399.

49. McColl TJ, Clarke DC. Kinetic modeling of leucine-mediated signaling and protein metabolism in human skeletal muscle. iScience. 2024;27(1):108634.

50. McCarthy RL, Zhang J, Zaret KS. Diverse heterochromatin states restricting cell identity and reprogramming. Trends in Biochemical Sciences. 2023;48(6):513–526.

51. Ugarte F, Sousae R, Cinquin B, Martin EW, Krietsch J, Sanchez G, Inman M, Tsang H, Warr M, Passegué E, Larabell CA, Forsberg EC. Progressive Chromatin Condensation and H3K9 Methylation Regulate the Differentiation of Embryonic and Hematopoietic Stem Cells. Stem Cell Reports. 2015;5(5):728–740.

52. May D, Yun S, Gonzalez DG, Park S, Chen Y, Lathrop E, Cai B, Xin T, Zhao H, Wang S, Gonzalez LE, Cockburn K, Greco V. Live imaging reveals chromatin compaction transitions and dynamic transcriptional bursting during stem cell differentiation in vivo. eLife. 2023;12:e83444.

53. Gentillon C, Li D, Duan M, Yu W-M, Preininger MK, Jha R, Rampoldi A, Saraf A, Gibson GC, Qu C-K, Brown LA, Xu C. Targeting HIF-1α in combination with PPARα activation and postnatal factors promotes the metabolic maturation of human induced pluripotent stem cell-derived cardiomyocytes. Journal of Molecular and Cellular Cardiology. 2019;132:120–135.

54. De Pater E, Ciampricotti M, Priller F, Veerkamp J, Strate I, Smith K, Lagendijk AK, Schilling TF, Herzog W, Abdelilah-Seyfried S, Hammerschmidt M, Bakkers J. Bmp Signaling Exerts Opposite Effects on Cardiac Differentiation. Circulation Research. 2012;110(4):578–587.

55. Tan J, Virtue S, Norris DM, Conway OJ, Yang M, Bidault G, Gribben C, Lugtu F, Kamzolas I, Krycer JR, Mills RJ, Liang L, Pereira C, Dale M, Shun-Shion AS, et al. Author Correction: Limited oxygen in standard cell culture alters metabolism and function of differentiated cells. The EMBO Journal. 2024;43(19):4439–4439.

56. Lygate CA. Maintaining energy provision in the heart: the creatine kinase system in ischaemia-reperfusion injury and chronic heart failure. Clinical Science (London, England: 1979). 2024;138(8):491–514.

57. Lopaschuk GD, Karwi QG, Tian R, Wende AR, Abel ED. Cardiac Energy Metabolism in Heart Failure. Circulation Research. 2021;128(10):1487–1513.

58. Zhong Y, Jia B, Xie C, Hu L, Liao Z, Liu W, Zhang Y, Huang G. Adenylate kinase 4 promotes neuronal energy metabolism and mitophagy in early cerebral ischemia via Parkin/PKM2 pathway. Experimental Neurology. 2024;377:114798.

59. Saddic LA, Nicoloro SM, Gupta OT, Czech MP, Gorham J, Shernan SK, Seidman CE, Seidman JG, Aranki SF, Body SC, Fitzgibbons TP, Muehlschlegel JD. Joint analysis of left ventricular expression and circulating plasma levels of Omentin after myocardial ischemia. Cardiovascular Diabetology. 2017;16(1):87.

60. Kitmitto A, Baudoin F, Cartwright EJ. Cardiomyocyte damage control in heart failure and the role of the sarcolemma. Journal of Muscle Research and Cell Motility. 2019;40(3–4):319–333.

61. Dias C, Nylandsted J. Plasma membrane integrity in health and disease: significance and therapeutic potential. Cell Discovery. 2021;7(1):4.

62. Vaniya A, Karlstaedt A, Gulkok D, Thottakara T, Liu Y, Fan S, Eades H, Vakrou S, Fukunaga R, Vernon HJ, Fiehn O, Abraham MR. Allele-specific dysregulation of lipid and energy metabolism in early-stage hypertrophic cardiomyopathy. Journal of Molecular and Cellular Cardiology Plus. 2024;8:100073.

63. Owen OE, Kalhan SC, Hanson RW. The Key Role of Anaplerosis and Cataplerosis for Citric Acid Cycle Function. Journal of Biological Chemistry. 2002;277(34):30409–30412.

64. Du C, Mukhi D, Li L, Li C, Pan S, Dumoulin B, Ha E, Kolligundla LP, Hou Y, Levinsohn J, Kang C, Klötzer KA, Wu J, Mohandes S, Wellen KE, et al. ACLY-Driven Metabolic Reprogramming Promotes Histone Acetylation and Inflammation-Associated Fibrosis in Chronic Kidney Disease. Advanced Science. 2026:e75247.

65. Meisner JK, Renberg A, Smith ED, Tsan Y-C, Elder B, Bullard A, Merritt OL, Zheng SL, Lakdawala NK, Owens AT, Ryan TD, Miller EM, Rossano JW, Lin KY, Claggett BL, et al. Low Penetrance Sarcomere Variants Contribute to Additive Risk in Hypertrophic Cardiomyopathy. Circulation. 2025;151(11):783–798.

66. García Hernandez S, de la Higuera Romero L, Fernandez A, Luisa Peña Peña M, Mora-Ayestaran N, Basurte-Elorz MT, Larrañaga-Moreira JM, Cárdenas Reyes I, Villacorta E, Valverde-Gómez M, Baustista-Paves A, Veira Villanueva E, Ortiz-Genga M, Lipov A, Brogger N, et al. Redefining the Genetic Architecture of Hypertrophic Cardiomyopathy: Role of Intermediate-Effect Variants. Circulation. 2025;152(15):1060–1075.

67. Miklas JW, Levy S, Hofsteen P, Mex DI, Clark E, Muster J, Robitaille AM, Sivaram G, Abell L, Goodson JM, Pranoto I, Madan A, Chin MT, Tian R, Murry CE, et al. Amino acid primed mTOR activity is essential for heart regeneration. iScience. 2022;25(1):103574.

68. Seeger T, Shrestha R, Lam CK, Chen C, McKeithan WL, Lau E, Wnorowski A, McMullen G, Greenhaw M, Lee J, Oikonomopoulos A, Lee S, Yang H, Mercola M, Wheeler M, et al. A Premature Termination Codon Mutation in MYBPC3 Causes Hypertrophic Cardiomyopathy via Chronic Activation of Nonsense-Mediated Decay. Circulation. 2019;139(6):799–811.

69. Prondzynski M, Krämer E, Laufer SD, Shibamiya A, Pless O, Flenner F, Müller OJ, Münch J, Redwood C, Hansen A, Patten M, Eschenhagen T, Mearini G, Carrier L. Evaluation of MYBPC3 trans-Splicing and Gene Replacement as Therapeutic Options in Human iPSC-Derived Cardiomyocytes. Molecular Therapy - Nucleic Acids. 2017;7:475–486.

70. Rust EM, Albayya FP, Metzger JM. Identification of a contractile deficit in adult cardiac myocytes expressing hypertrophic cardiomyopathy–associated mutant troponin T proteins. Journal of Clinical Investigation. 1999;103(10):1459–1467.

71. Coppini R, Ho CY, Ashley E, Day S, Ferrantini C, Girolami F, Tomberli B, Bardi S, Torricelli F, Cecchi F, Mugelli A, Poggesi C, Tardiff J, Olivotto I. Clinical Phenotype and Outcome of Hypertrophic Cardiomyopathy Associated With Thin-Filament Gene Mutations. Journal of the American College of Cardiology. 2014;64(24):2589–2600.

72. Pioner JM, Vitale G, Gentile F, Scellini B, Piroddi N, Cerbai E, Olivotto I, Tardiff J, Coppini R, Tesi C, Poggesi C, Ferrantini C. Genotype-Driven Pathogenesis of Atrial Fibrillation in Hypertrophic Cardiomyopathy: The Case of Different TNNT2 Mutations. Frontiers in Physiology. 2022;13:864547.

73. Gilbert CJ, Longenecker JZ, Accornero F. ERK1/2: An Integrator of Signals That Alters Cardiac Homeostasis and Growth. Biology. 2021;10(4):346.

74. Davis J, Davis LC, Correll RN, Makarewich CA, Schwanekamp JA, Moussavi-Harami F, Wang D, York AJ, Wu H, Houser SR, Seidman CE, Seidman JG, Regnier M, Metzger JM, Wu JC, et al. A Tension-Based Model Distinguishes Hypertrophic versus Dilated Cardiomyopathy. Cell. 2016;165(5):1147–1159.

75. Pettinato AM, Ladha FA, Mellert DJ, Legere N, Cohn R, Romano R, Thakar K, Chen Y-S, Hinson JT. Development of a Cardiac Sarcomere Functional Genomics Platform to Enable Scalable Interrogation of Human *TNNT2* Variants. Circulation. 2020;142(23):2262–2275.

76. Zhou R, Li J, Zhang L, Cheng Y, Yan J, Sun Y, Wang J, Jiang H. Role of Parkin-mediated mitophagy in glucocorticoid-induced cardiomyocyte maturation. Life Sciences. 2020;255:117817.

77. Yang J, Ding N, Zhao D, Yu Y, Shao C, Ni X, Zhao Z-A, Li Z, Chen J, Ying Z, Yu M, Lei W, Hu S. Intermittent Starvation Promotes Maturation of Human Embryonic Stem Cell-Derived Cardiomyocytes. Frontiers in Cell and Developmental Biology. 2021;9:687769.

