## Supplementary figures for "Molecular Rewiring in TNNT2-Linked Hypertrophic Cardiomyopathy hiPSC-Derived Cardiomyocytes upon Metabolic Maturation"

### Supplementary Information

#### Supplementary methods

##### Seahorse XF Mito Stress Test

Seahorse metabolic profiling was performed approximately 3 days after replating hiPSC-derived cardiomyocytes at a density of 75,000 cells per well on Matrigel-coated Seahorse XFe96 cell culture plates. One hour prior to the assay, cells were switched to Mito Stress Test assay medium consisting of RPMI 1640 lacking sodium bicarbonate and glucose (Sigma-Aldrich, #R1383), supplemented with 10 mM glucose (Sigma-Aldrich, #71718), 2 mM glutamine (Sigma-Aldrich, #G8540), and 1 mM sodium pyruvate (Sigma-Aldrich, #P5280). Plates were incubated in a non-$CO_2$ incubator for up to 50 minutes before initiating the experiment. Oxygen consumption rate (OCR) was measured using the Agilent Seahorse XFe96 analyzer. The Mito Stress Test was conducted by sequential injection of oligomycin (1 µM; Sigma-Aldrich, Cat. No. #04876), FCCP (1.5 µM; Sigma-Aldrich, # C2920), and the mitochondrial inhibitors rotenone (100 nM; Sigma-Aldrich, #557368) and antimycin A (1 µM; Sigma-Aldrich, #A8674). The Fatty acid oxidation (FAO) Stress Test was performed following a protocol similar to the Mito Stress Test, but with the additional injection of 4 µM Etomoxir (Sigma, 236020) to inhibit CPT1-dependent FAO, or Seahorse assay medium as a control in non-inhibited cells. Following the experiment, cell numbers were determined, and raw data were normalized based on cell numbers per well in Wave software (Agilent Technologies, Inc.) and visualized using GraphPad Prism software (version 9.5.1).

##### Cloning

The mTORC1 biosensor-encoding plasmid (Addgene ID: 140828, 9) was obtained to study mTORC1 signaling in live cells. Plasmid extraction was performed, followed by PCR-based cloning of the biosensor-encoding sequence into the lentiviral transfer plasmid pHIV-EGFP (Addgene ID: 21373), with omission of the EGFP-encoding sequence that overlaps with the biosensor YPET spectra. Plasmid extraction was performed from the modified pHIV-EGFP plasmids, and the samples were submitted for Sanger sequencing.

##### Generation of a stable hiPSC AIMTOR biosensor line

HEK293FT cells were seeded in T75 flasks for mass virus production. The cells were co-transfected with four Addgene plasmids (3rd generation lentivirus): the AIMTOR-encoding pHIV-EGFP transfer plasmid, the envelope plasmid (pMD2.G, ID: 12259), and two packaging plasmids (pRSV-Rev, ID: 12253; pMDLg, ID: 12251). Lipofectamine 3000 (Invitrogen, #L3000001) was used for overnight transfection. The cell culture supernatant containing the produced lentivirus was collected ~48–72 hours after transfection. The viruses were concentrated using ultracentrifugation (25,000 × g, 4 °C, 2 hours). The qPCR Lentivirus Titer Kit (Applied Biological Materials, LV900) was used for virus titration. To generate a stable hiPSC AIMTOR biosensor line, hiPSC medium was refreshed ~16 hours after transduction with lentivirus at a multiplicity of infection (MOI) of 1. Fluorescence-Activated Cell Sorting (FACS) was employed to isolate $YFP^+$ cells with successful expression of the AIMTOR biosensor. The sorted hiPSCs were seeded at a density of 4,000 cells per well in ClonR medium (StemCell Technologies,#05888) for up to 4 days until the appearance of single individual colonies. The colonies were subsequently manually picked for expansion and cryopreservation.

**Bioluminescence resonance energy transfer (BRET)**

The Day 16 hiPSC-CMs, after metabolic selection, were replated into Nunc™ F96 MicroWell™ White Polystyrene Plates (Thermo Scientific™, #1256602) coated with Matrigel at a density of 75,000 cells per well. The hiPSC-CMs were subjected to metabolic maturation treatment for 4 weeks. The Mithras LB 940 instrument (Berthold Technologies) was used to record luminescence and fluorescence emissions. Recordings were performed after the addition of 2 μM luciferase substrate Coelenterazine H (NanoLight Technology, #301) in the dark, followed by 5 minutes of incubation. BRET ratios were obtained based on the emission signal ratio of 530 nm to 480 nm. The hiPSC-CM medium was changed to RPMI without phenol red (Sigma, #R8755) to prevent interference with assay measurements.

**CUTAC Data analysis**

The sequencing reads were first aligned to the human genome (hg38) using Bowtie2. In parallel, sample reads were aligned to the *Drosophila* BDGP6 genome build to infer the percentage of reads for spike-in–based normalization. The sorted BAM files were then converted to BigWig (BW) files based on normalization scale factors. Replicates were merged using BigWigMerge. Peak calling was performed with IgG as the control using SEACR[78] stringent settings and a false discovery rate (FDR) threshold of <0.001. Peaks were visualized in the Integrative Genomics Viewer (IGV). EdgeR[79] was used for differential chromatin accessibility analysis, and HOMER[80] motif enrichment analysis was performed to identify the presence of known transcription factor binding motifs. The GREAT genomic tool[81] was used to assess the association of accessible regions with genes and potential biological functions using the “Basal plus extension” configuration (5 kb upstream, 1 kb downstream, and up to 1000 kb distal). All heatmaps and plots related to chromatin accessibility data were generated using R and the DeepTools suite plotting functions.

**Protein expression analysis**

hiPSC-CMs (Day 30 and 44 of differentiation) were centrifuged at 200 × g for 7 minutes, and the cell pellets were resuspended in lysis buffer (100 mM HEPES, pH 8–8.5 (Sigma, #H3375); and 20% SDS,(Sigma-Aldrich, #L4509)) with a volume of 11.25 µL per 300,000 hiPSC-CMs.The lysate was processed as follows: heating at 95 °C for 5 minutes, cooling on ice for 3 minutes, and two rounds of sonication for 30 seconds each. Benzonase (EMD Millipore-Sigma, #70664) treatment at 37 °C for 30 minutes was used for chromatin fragmentation, followed by lysate reduction using 10 mM dithiothreitol (DTT; Thermo Fisher Scientific, #R0861) at 37 °C for 30 minutes. Subsequently, 50 mM chloroacetamide (CAA; Sigma-Aldrich, #C0267) was applied for 30 minutes in the dark to alkylate the lysates. Then, 50 mM DTT was applied for 10 minutes to quench the reaction.

SP3 technology (Single-Pot Solid-Phase-enhanced Sample Preparation) was employed with Sera-Mag Speed Beads (GE Life Sciences, #05855) in a 1:1 ratio. Protein binding to the beads was carried out in 80 % ethanol, followed by two washes with 90 % ethanol. Overnight digestion with trypsin (Thermo Fisher Scientific, #17504044, 1:50 w/w) was then performed.

The processed samples were eluted using BioPure midi-columns (Nest Group Inc., #05888) with methanol and sequential washes including trifluoroacetic acid (TFA; Thermo Fisher Scientific, #A12198.22), formic acid (FA; Thermo Fisher Scientific, #28905), and acetonitrile (ACN; Thermo Fisher Scientific, #047138.K2). The pH was adjusted to 3–4, and final elution was conducted with 0.1 % FA in 60 % ACN. Sample concentration and protein quantification were performed using SpeedVac concentrator (Thermo Fisher Scientific) and NanoDrop spectrophotometer (Thermo Fisher Scientific). The eluates were resuspended in 0.1 % FA for MS analysis. Proteomic analysis was performed on a Q Exactive HF Orbitrap mass spectrometer coupled with an Easy-nLC liquid chromatography system (Thermo Scientific). MaxQuant (version 2.0.0.0) and Proteome Discoverer (Thermo Fisher Scientific), with a false discovery rate set at 1%, were used for data processing. The human proteome database (UniProt, release 2022_06) was used, excluding “Potential contaminant/REV” peptides. Final raw data analysis, including log2 transformation and normalization of protein intensities, was performed using Perseus software. R (version 4.2.3) was employed for visualization and analysis.

**The integration of OMICS datasets**

To understand the overlap between mRNA and protein expression, a threshold of $|\log_2(\text{fold change})| \geq 1$ and adjusted $p < 0.05$ was used to distinguish differentially expressed genes from the RNA-seq and proteomics datasets. The VennDiagram package in R was used to identify overlapping gene sets for enrichment analysis. Differentially accessible regions were annotated using the HOMER annotation tool[82], and the UpSetPlot Python package was used to identify genes showing different patterns of expression.

#### Supplementary Figures and Legends

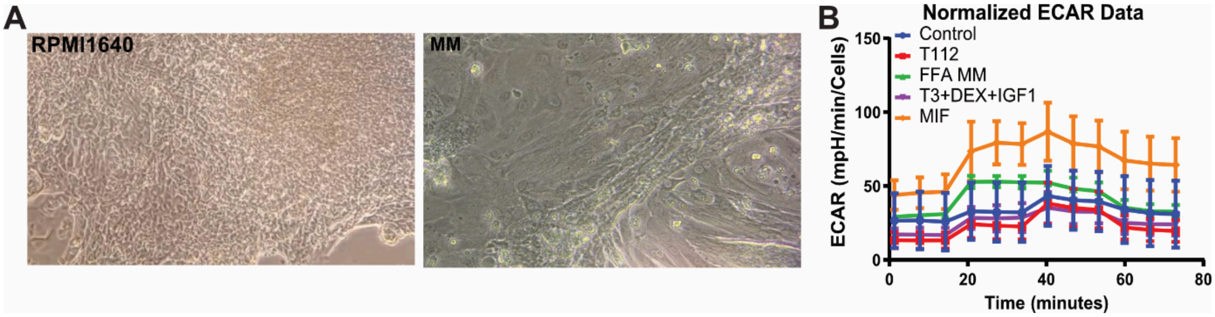


**Supplementary Figure 1 (related to Figure 1). Metabolic profiling of MIF-treated hiPSC-CMs using Seahorse Mito Stress Test analysis.**

**(A)** Representative brightfield images captured at 20× objective of hiPSC-CMs cultured in standard RPMI1640 medium or metabolically matured (MM) conditions. **(B)** Extracellular acidification rate (ECAR) kinetics normalized to cell number.

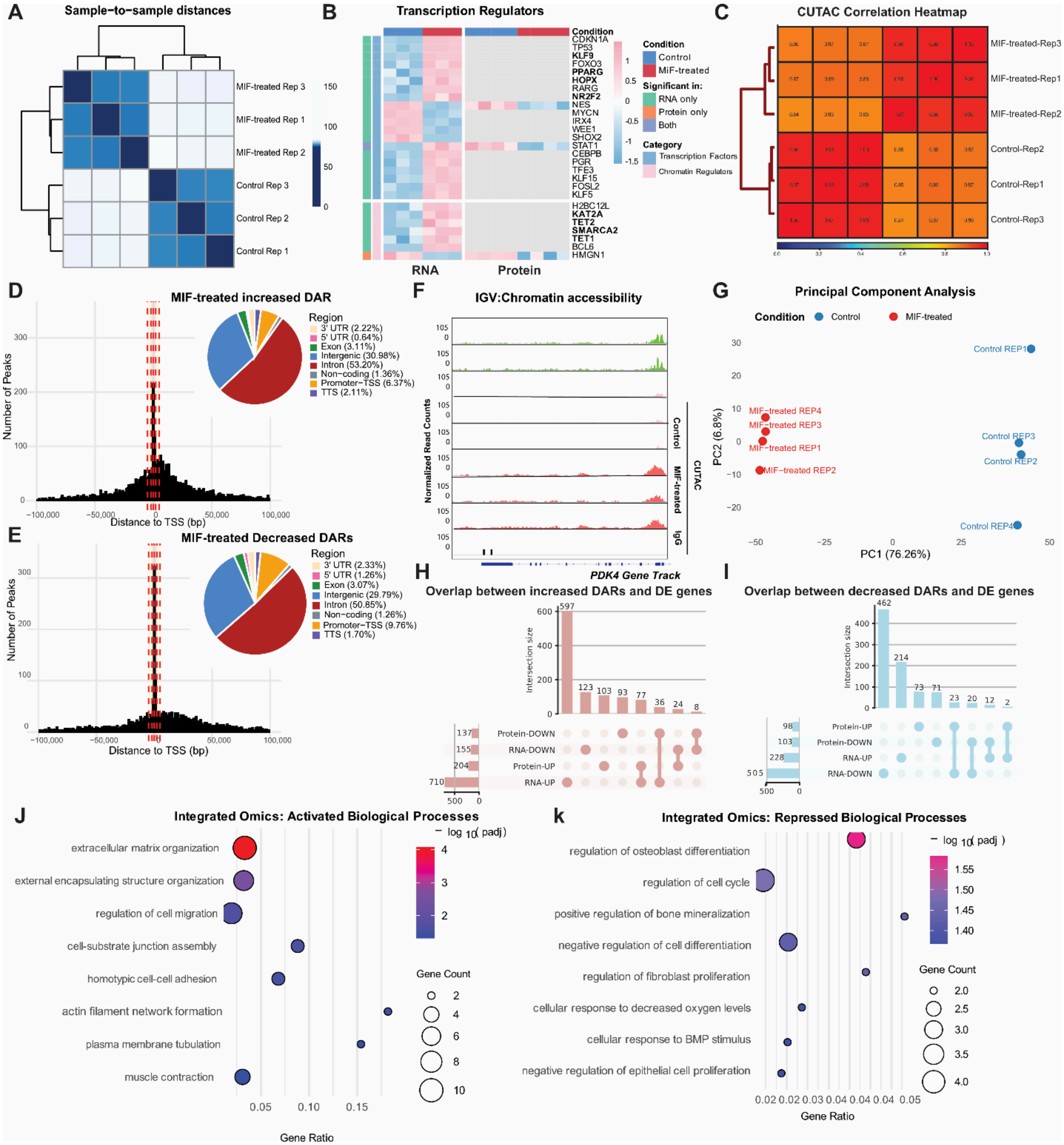


**Supplementary Figure 2 (related to Figure 2). Additional Multi-omics characterization of MIF-treated hiPSC-CMs**.

**(A)** Sample-to-sample distance heatmap of RNA-seq datasets from control and MIF-treated hiPSC-CMs. **(B)** Heatmap showing selected transcription factors and chromatin regulators identified by integrated RNA-seq and proteomics analyses. **(C)** Sample correlation heatmap of CUTAC datasets. **(D)** Distance-to-transcription start site (TSS) histogram and genomic annotation of regions with increased chromatin accessibility in MIF-treated hiPSC-CMs. **(E)** Distance-to-TSS histogram and genomic annotation of regions with decreased chromatin accessibility in MIF-treated hiPSC-CMs. **(F)** Representative CUTAC genome browser tracks showing chromatin accessibility at the *PDK4* locus. **(G)** Principal component analysis (PCA) of Proteomics datasets from control and MIF-treated hiPSC-CMs. **(H–I)** Overlap between differentially accessible region (DAR)-associated genes and differentially expressed genes (DEGs) at the RNA and protein levels for regions with increased (H) or decreased (I) accessibility. **(J–K)** Gene Ontology enrichment analysis of activated (J) and repressed (K) biological processes identified by integrated multi-omics analysis. Statistical significance: adjusted $p \leq 0.05$ unless otherwise indicated ($n$ = 3–4 independent differentiations).

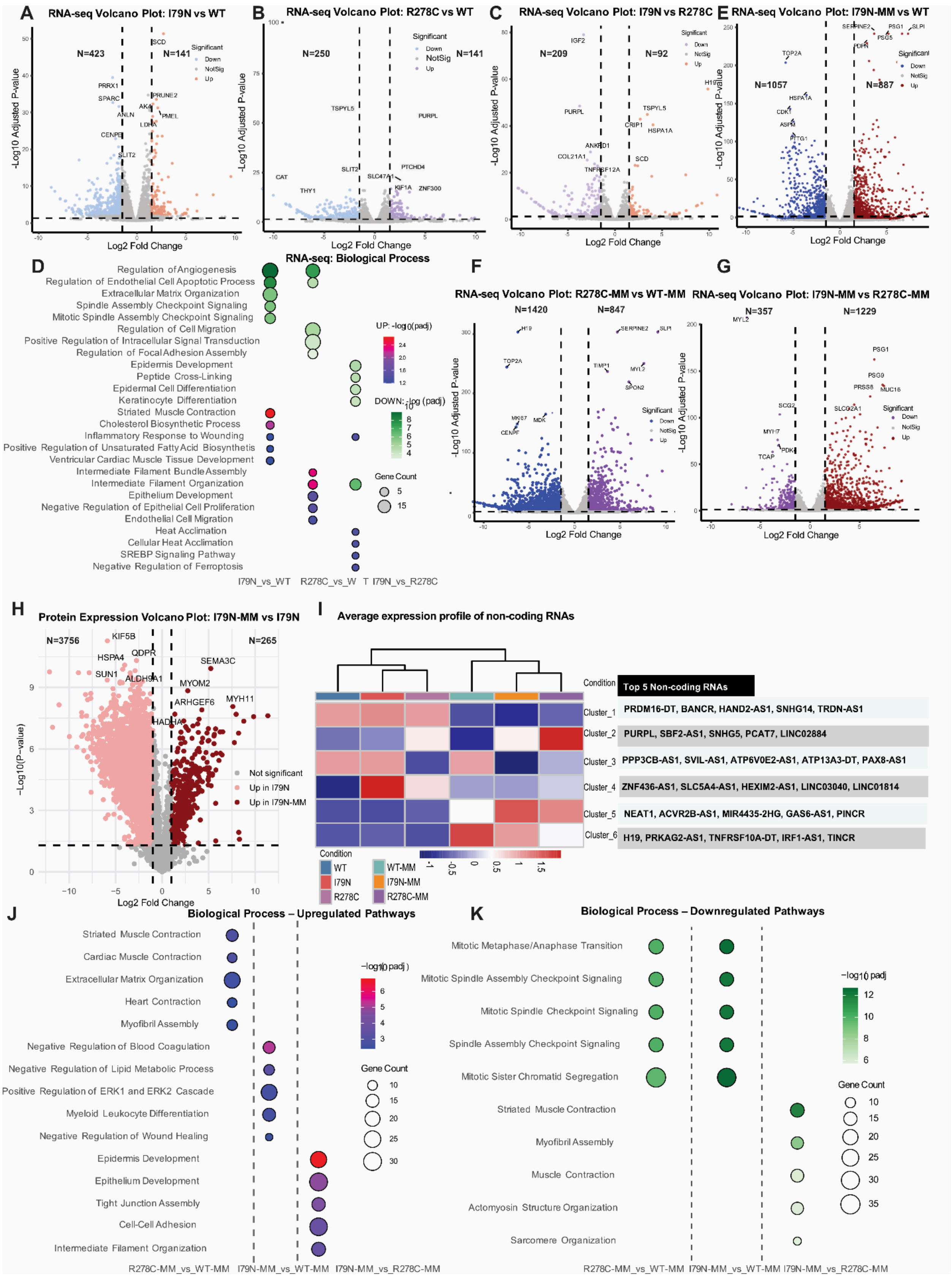
A
RNA-seq Volcano Plot: I79N vs WT
B
RNA-seq Volcano Plot: R278C vs WT
C
RNA-seq Volcano Plot: I79N vs R278C
E
RNA-seq Volcano Plot: I79N-MM vs WT
D
RNA-seq: Biological Process
F
RNA-seq Volcano Plot: R278C-MM vs WT-MM
G
RNA-seq Volcano Plot: I79N-MM vs R278C-MM
H
Protein Expression Volcano Plot: I79N-MM vs I79N
I
Average expression profile of non-coding RNAs
Top 5 Non-coding RNAs
Cluster_1 PRDM16-DT, BANCR, HAND2-AS1, SNHG14, TRDN-AS1
Cluster_2 PURPL, SBF2-AS1, SNHG5, PCAT7, LINC02884
Cluster_3 PPP3CB-AS1, SVIL-AS1, ATP6V0E2-AS1, ATP13A3-DT, PAX8-AS1
Cluster_4 ZNF436-AS1, SLC5A4-AS1, HEXIM2-AS1, LINC03040, LINC01814
Cluster_5 NEAT1, ACVR2B-AS1, MIR4435-2HG, GAS6-AS1, PINCR
Cluster_6 H19, PRKAG2-AS1, TNFRSF10A-DT, IRF1-AS1, TINCR
J
Biological Process – Upregulated Pathways
K
Biological Process – Downregulated Pathways

**Supplementary Figure 3 (related to Figure 4). Additional transcriptomic and proteomic analyses of TNNT2-linked HCM hiPSC-CMs.**

**(A–C)** RNA-seq volcano plots showing differentially expressed genes between I79N vs WT (A), R278C vs WT (B), I79N vs R278C (C) **(D)** Gene Ontology biological process enrichment analysis of differentially expressed genes across indicated comparisons using RNA-seq dataset. **(E–G)** RNA-seq volcano plots showing differentially expressed genes between I79N-MM vs WT-MM (E), R278C-MM vs WT-MM (F), and I79N-MM vs R278C-MM (G). **(H)** Proteomics volcano plot showing differentially expressed proteins between I79N-MM and I79N hiPSC-CMs. **(I)** Heatmap showing expression patterns of selected non-coding RNAs across experimental groups. The top five non-coding RNAs from each cluster are indicated. **(J–K)** Gene Ontology biological process enrichment analysis of upregulated (J) and downregulated (K) pathways identified from RNA-seq datasets. Statistical significance: adjusted $p \leq 0.05$ ($n$ = 3-4 independent differentiations for RNA-seq and proteomics).

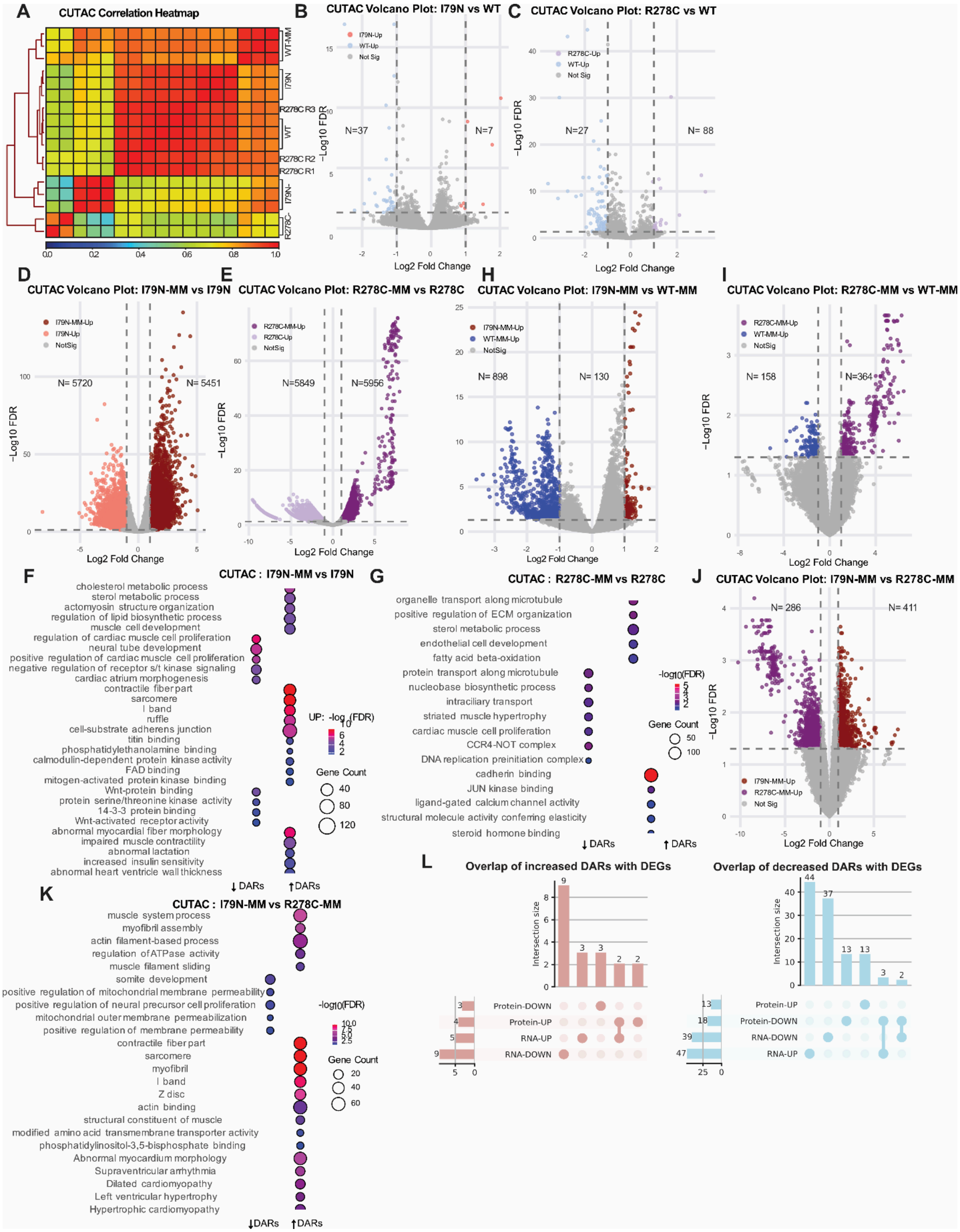

**Supplementary Figure 4 (related to Figure 5). Additional chromatin accessibility analyses of TNNT2-linked HCM hiPSC-CMs.**

**(A)** CUTAC sample correlation heatmap across WT, I79N, and R278C hiPSC-CMs under immature and metabolically matured (MM) conditions. **(B–C)** CUTAC volcano plots showing differentially accessible regions (DARs) between I79N vs WT (B), R278C vs WT (C). **(D–G)** CUTAC volcano plots showing DARs between I79N-MM vs I79N(D) and R278C-MM vs R278C (E). Gene Ontology enrichment analysis of DAR-associated genes identified in I79N-MM vs I79N (F) and R278C-MM vs R278C (G). **(H–I)** CUTAC volcano plots showing differentially accessible regions between I79N-MM vs WT-MM (H) and R278C-MM vs WT-MM (I). **(J)** CUTAC volcano plot showing differentially accessible regions between I79N-MM and R278C-MM hiPSC-CMs. **(K)** Gene Ontology enrichment analysis of DAR-associated genes identified in I79N-MM vs R278C-MM. **(L)** Overlap between DAR-associated genes and differentially expressed genes (DEGs) at the RNA and protein levels for regions with increased or decreased chromatin accessibility. Statistical significance: adjusted $p \leq 0.05$ unless otherwise indicated ($n$ = 3 independent differentiations).

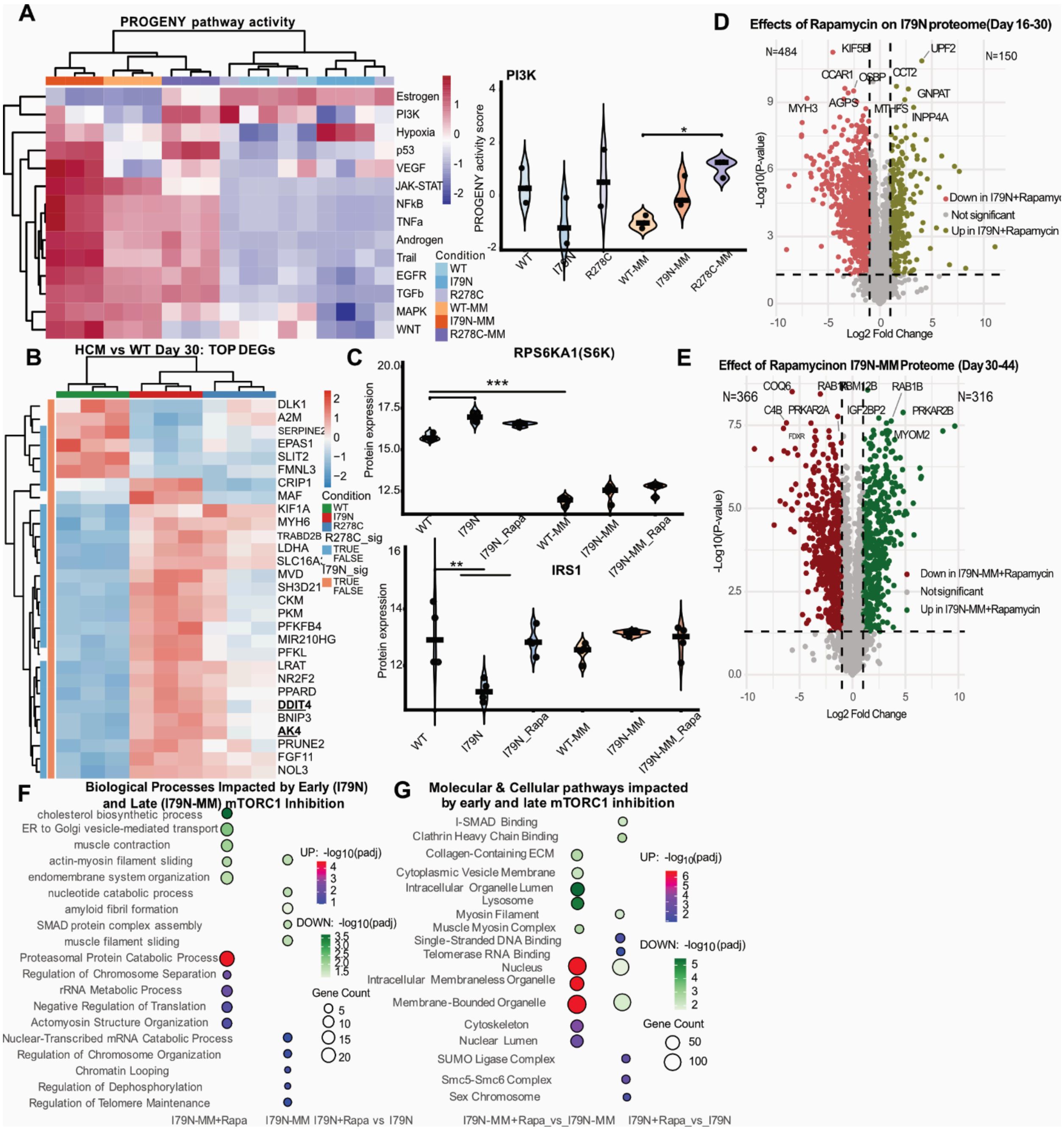


**Supplementary Figure 5 (related to Figure 6). Additional analyses of the effects of mTORC1 inhibition on TNNT2-linked HCM hiPSC-CMs during metabolic maturation.**

**(A)** PROGENy pathway activity analysis across WT, I79N, and R278C hiPSC-CMs under immature and metabolically matured (MM) conditions. Violin plots show PI3K pathway activity scores. **(B)** Heatmap of Top differentially expressed genes associated with HCM pathology in WT and TNNT2 variant hiPSC-CMs at day 30. **(C)** Protein expression levels of RPS6KA1 (S6K) and IRS1 across experimental groups determined by proteomic analysis. **(D–E)** Proteomic volcano plots showing differentially expressed proteins following rapamycin treatment in I79N hiPSC-CMs during the immature stage (D; I79N+Rapa vs I79N) and following metabolic maturation (E; I79N-MM+Rapa vs I79N-MM). **(F)** Gene Ontology biological process enrichment analysis of proteins altered by early and late mTORC1 inhibition. **(G)** Molecular and cellular pathways affected by rapamycin treatment identified through Gene Ontology enrichment analysis. Statistical significance: adjusted $p \leq 0.05$ ($n$ = 3 independent differentiations). $p \leq 0.05$, **p** ≤ 0.01, ***p*** ≤ 0.001, **p** ≤ 0.0001.
